# A conserved population of genetically defined striatal neurons gates opioid reward

**DOI:** 10.64898/2026.07.28.741269

**Authors:** Olivia R. Drake, Caroline M. Fiore, Emily T. Jorgensen, Catherine E. Newman, Luke A. Potter, Austyn Trull, Aditya Pradeep, Lara Ianov, Jamie Peters, Jasper A. Heinsbroek, Jeremy J. Day

## Abstract

A longstanding paradox in striatal circuit architecture is that opioid reward depends on μ-opioid receptors (µORs) in nucleus accumbens medium spiny neurons (MSNs), yet µOR function is not explained by the canonical D1/direct and D2/indirect pathway organization. Here, we identify a rare MSN population marked by *Chst9* that exhibits exceptionally high expression of the µOR and is conserved across species. Notably, Chst9-MSNs comprise a specialized indirect pathway striatal neuron subtype that is molecularly and spatially distinct from canonical striatal populations. Opioids robustly silence Chst9-MSNs, and selective deletion of *Oprm1* from this population abolishes fentanyl-conditioned place preference. These findings establish Chst9-MSNs as a critical substrate for opioid reward and define a new cellular framework for therapies targeting opioid use disorder.

## INTRODUCTION

Opioids reinforce behavior primarily through activation of mu opioid receptors (µORs), encoded by *Oprm1*, within mesolimbic reward circuitry^1^. Opioid reward is often attributed to µOR-mediated disinhibition of dopaminergic neurons in the ventral tegmental area (VTA)^2–4^. However, µORs are also expressed in the nucleus accumbens (NAc), suggesting that local accumbal mechanisms contribute to opioid reinforcement^2,5–7^. Consistent with this idea, direct infusion of µOR agonists into the NAc is reinforcing^8–12^, selective restoration of µORs in a subset of striatal neurons rescues opioid reward following global *Oprm1* deletion^13^, and targeted ablation of *Oprm1*-expressing cells in the NAc disrupts heroin seeking^14^. Together, these findings establish the NAc as a critical site of µOR-dependent opioid reward, yet how local µOR signaling is organized within canonical striatal circuitry remains unclear.

The principal neurons of the NAc are medium spiny neurons (MSNs), traditionally classified as D1-MSN or D2-MSN subtypes based on dopamine receptor expression^15–19^. Recent single nucleus RNA sequencing (snRNA-seq) studies, however, have revealed extensive molecular diversity beyond this D1/D2 framework^20–26^. Notably, *Oprm1* expression is unevenly distributed across MSN populations and is present in subsets of both D1- and D2-MSNs^22,23,27^. Thus, although accumbal µOR signaling is critical for opioid reward, its cellular organization is not readily explained by the canonical D1/direct and D2/indirect pathway framework, raising the possibility that opioid reward is mediated by specialized, molecularly defined MSN subpopulations.

Here, we combine single cell transcriptomics, high resolution spatial profiling, circuit mapping, electrophysiology, and cell type specific genetic manipulation to define the molecular and functional properties of a sparse MSN population marked by the carbohydrate sulfotransferase 9 gene (*Chst9*; termed Chst9-MSNs). We identify Chst9-MSNs as a transcriptionally distinct, µOR-enriched MSN subtype that is conserved across species, forms discrete spatial clusters along the border of the NAc, and preferentially projects to the ventral pallidum (VP). Chst9-MSNs are directly inhibited by µOR activation, and selective deletion of *Oprm1* from these neurons abolishes fentanyl reward learning. Together, these findings identify this specialized, non-canonical MSN population as a critical cellular substrate for opioid reward learning and provide a cellular framework for understanding how accumbal µOR signaling is organized beyond canonical D1/D2 classifications.

## RESULTS

### Chst9-MSNs are a transcriptionally distinct, Oprm1-enriched MSN subtype

To examine *Oprm1* expression across cell types in the NAc, we performed snRNA-seq on 44,592 nuclei collected from 6 male and 6 female Sprague-Dawley rats. Dimensionality reduction and clustering of the final integrated object resulted in 19 transcriptionally distinct populations (**Fig. 1a**), including known *Drd1*-expressing and *Drd2*-expressing MSNs, somatostatin, parvalbumin, and cholinergic interneurons, and non-neurons previously identified in the NAc^21,22^. Cell types were annotated using established marker genes and validated using the Allen Institute’s MapMyCells tool^28^ (**Fig. 1a**).

**Figure 1.**
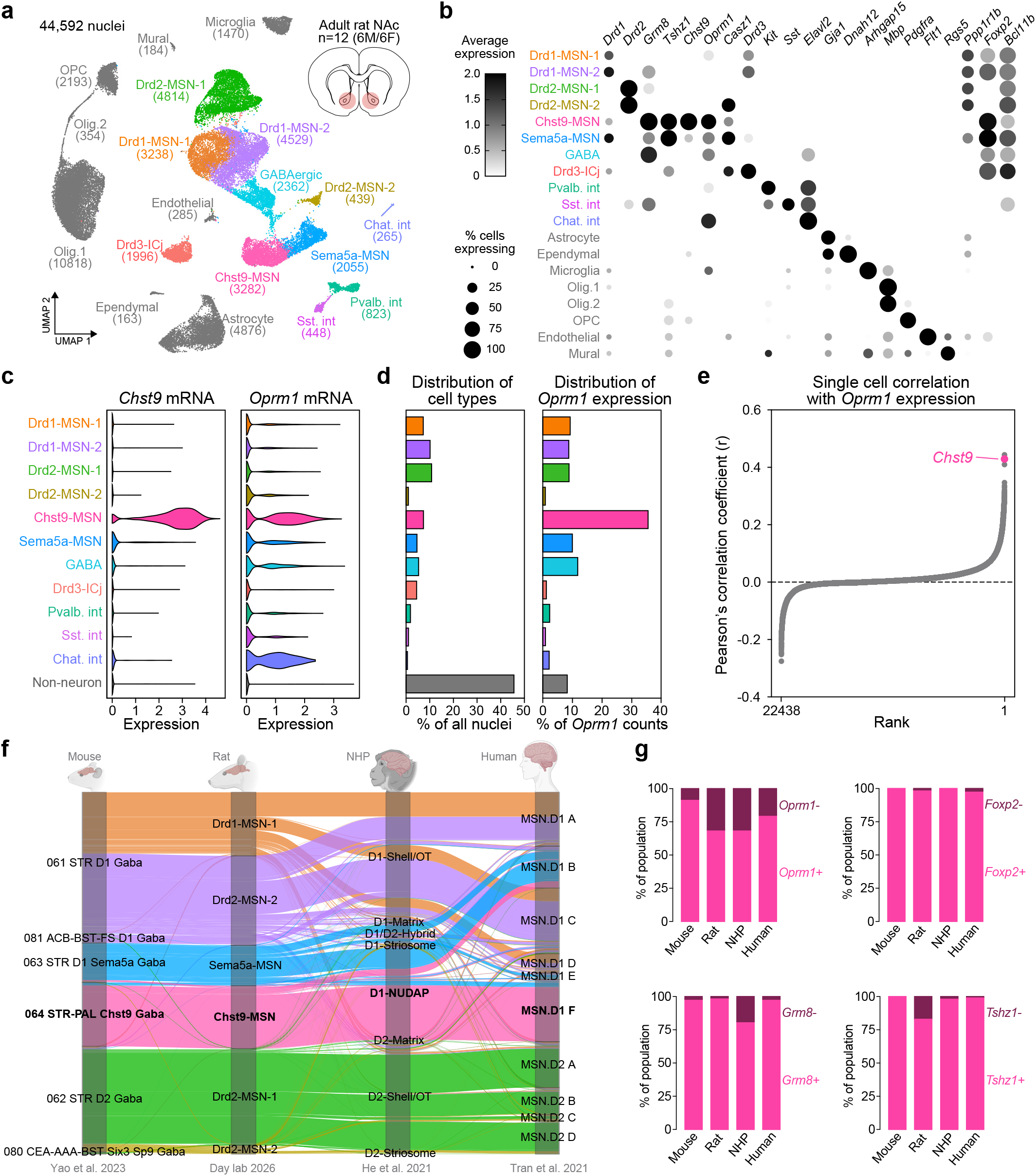
snRNA-seq identifies Chst9-MSNs as a transcriptionally distinct, µOR-enriched MSN subtype that is conserved across species. **a**, NAc tissue from male and female rats (n = 6 per sex) was processed for snRNA-seq using the 10x Genomics platform. Clustering of 44,592 nuclei identifies major rat NAc cell classes, including *Chst9*-expressing MSNs (Chst9-MSNs). **b**, Dot plot showing expression and proportion of cells expressing marker genes across cell types. **c**, Violin plots showing *Chst9* and *Oprm1* expression across cell types. **d**, Distribution of cell types and proportion of *Oprm1* transcripts contributed by each cell type. **e**, Ranked scatter plot showing single cell correlation of each detected gene with *Oprm1* across all annotated cell populations. **f**, Hierarchical correlation mapping across mouse (Yao et al., 2023), rat (this study), NHP (He et al., 2021), and human (Tran et al., 2021) datasets. Rat Chst9-MSNs correspond to STR-PAL Chst9 Gaba neurons in mouse, D1-NUDAP neurons in NHP, and D1.MSN-F neurons in human. **g**, The proportion of *Oprm1*+, *Foxp2*+, *Grm8*+, and *Tshz1*+ cells within homologous Chst9-MSN populations.

Previously, we identified *Grm8*-expressing MSNs as a subtype of D1-MSNs^21^, and subsequent work demonstrated that *Chst9* marks a subset of Grm8-MSNs enriched for *Oprm1*^*29*^. Analysis of the present dataset revealed that Grm8-MSNs are composed of two transcriptionally distinct populations, termed Chst9-MSNs and Sema5a-MSNs based on subclass annotation in the Allen Institute’s Mouse Whole-Brain Transcriptomic Cell Type Atlas reference taxonomy^30^. Both populations express canonical MSN markers including *Bcl11b* and *Foxp2* while Chst9-MSNs are distinguished by selective enrichment for *Chst9* and *Oprm1* (**Fig. 1b, c** and **S1**). In contrast, *Sema5a* expression was not restricted to Sema5a-MSNs, and this population lacks a uniquely selective marker. Instead, Sema5a-MSNs share transcriptional features with both Drd2-MSN-2 and Chst9-MSN populations.

For example, differential expression analysis revealed that Drd2-MSN-2 and Sema5a-MSNs both express *Casz1* and *Alk*, which have been described in non-canonical “eccentric MSN”^31^ and “D1/D2 hybrid”^24^ populations (**Fig. 1b** and **S1**). Chst9-MSNs and Sema5a-MSNs overlap in their expression of genes associated with these MSN populations, including *Otof, Adarb2*, and *Ppm1e* (**Fig. S1**)^24,31,32^. Additionally, *Tshz1* is expressed across both Chst9-MSNs and Sema5a-MSNs. The neuronal population expressing this transcription factor has been implicated in reward- and aversion-related circuitry^33^ and is located in cell clusters along the NAc boundary^20^. *Tshz1* was also recently identified as a marker of µOR-enriched MSN populations involved in opioid withdrawal aversion^34^. Despite these shared features, Chst9-MSNs and Sema5a-MSNs exhibit distinct transcriptional profiles. Chst9-MSNs differentially express *Vwc2l* and *Col14a1*, markers previously associated with a subtype of D1 striosomal MSN^32^. Although Chst9-MSN and Sema5a-MSN populations can be distinguished by differential gene expression, many of these transcripts exhibit graded expression across these MSN subtypes, consistent with their close transcriptional relationship (**Fig. S1b**).

Importantly, *Chst9* selectively identifies the MSN population with the highest *Oprm1* expression, indicating that targeting this gene may allow selective access to this subtype (**Fig. 1c**). Indeed, although Chst9-MSNs comprise only ∼7% of the total NAc cells, they contain over one third (35.3%) of all *Oprm1* transcripts detected in the region (**Fig. 1d**). Consistent with this enrichment, ranked single-cell correlation analysis identified *Chst9* among the top transcripts associated with *Oprm1* expression across NAc cell populations (**Fig. 1e**), supporting its use as a marker of this *Oprm1*-enriched MSN subtype. Together, these findings identify Chst9-MSNs as a transcriptionally distinct MSN subtype that is disproportionately enriched for *Oprm1*.

### Chst9-MSNs represent a conserved Oprm1-enriched MSN subtype across mammalian species

While Chst9-MSNs share transcriptional features with MSN populations identified in other species^24,31^, we next assessed their transcriptional correspondence to previously defined NAc MSN subtypes. To do this, we performed cross-species cell type mapping using the Allen Institute’s MapMyCells framework^28^ to compare rat MSN populations to published snRNA-seq datasets from mouse^30^, non-human primate (NHP)^24^, and human^26^ striatal tissue. Rat MSN clusters (Drd1-MSN-1, Drd1-MSN-2, Drd2-MSN-1, Drd2-MSN-2, Chst9-MSN, and Sema5a-MSN) were mapped independently to each species-specific reference taxonomy following ortholog conversion of one-to-one gene identifiers and restriction to conserved orthologous genes. Across species, Chst9-MSNs consistently mapped most strongly to specific MSN subtypes previously annotated in each reference dataset, including STR-PAL Chst9 Gaba neurons in mouse, D1-NUDAP neurons in NHP, and D1.MSN-F neurons in human (**Fig. 1f**). These observations align with recent spatiomolecular profiling of the human NAc, which identified a transcriptionally distinct DRD1-MSN population enriched for *OPRM1* that shares transcriptional features with Chst9-MSNs, organized in spatially clustered “islands” along the perimeter of the NAc shell subregion^35^.

Given that elevated *Oprm1* expression is a defining feature of rat Chst9-MSNs, with 69.2% expressing *Oprm1*, wenextassessedwhetherthis *Oprm1* enrichment was conserved across correspondingly mapped MSN populations in other species. Notably, homologous MSN populations exhibited elevated proportions of *Oprm1*-expressing cells in mouse (92.0%), NHP (68.8.0%), and human (80.2%) datasets (**Fig. 1g**), demonstrating conservation of *Oprm1* enrichment among transcriptionally corresponding Chst9-MSN populations. Additional Chst9-MSN marker genes including *Grm8, Foxp2*, and *Tshz1* also showed conserved expression across species, further supporting preservation of Chst9-MSN molecular identity.

### Chst9-MSNs occupy a discrete spatial domain along the NAc shell border

To investigate the spatial organization of Chst9-MSNs, we performed high resolution, single cell spatial transcriptomics with the Xenium platform (10x Genomics, **Fig. 2a**). Four coronal sections spanning anterior-posterior coordinates between Bregma +2.04 and +0.60 mm were collected from each of four adult naive C57BL/6 mice (2 male, 2 female). Tissue sections were processed using the standard Xenium mouse probe panel supplemented with 100 custom probes selected to stratify NAc neuronal populations (**Fig. 2a, Table S1**).

**Figure 2.**
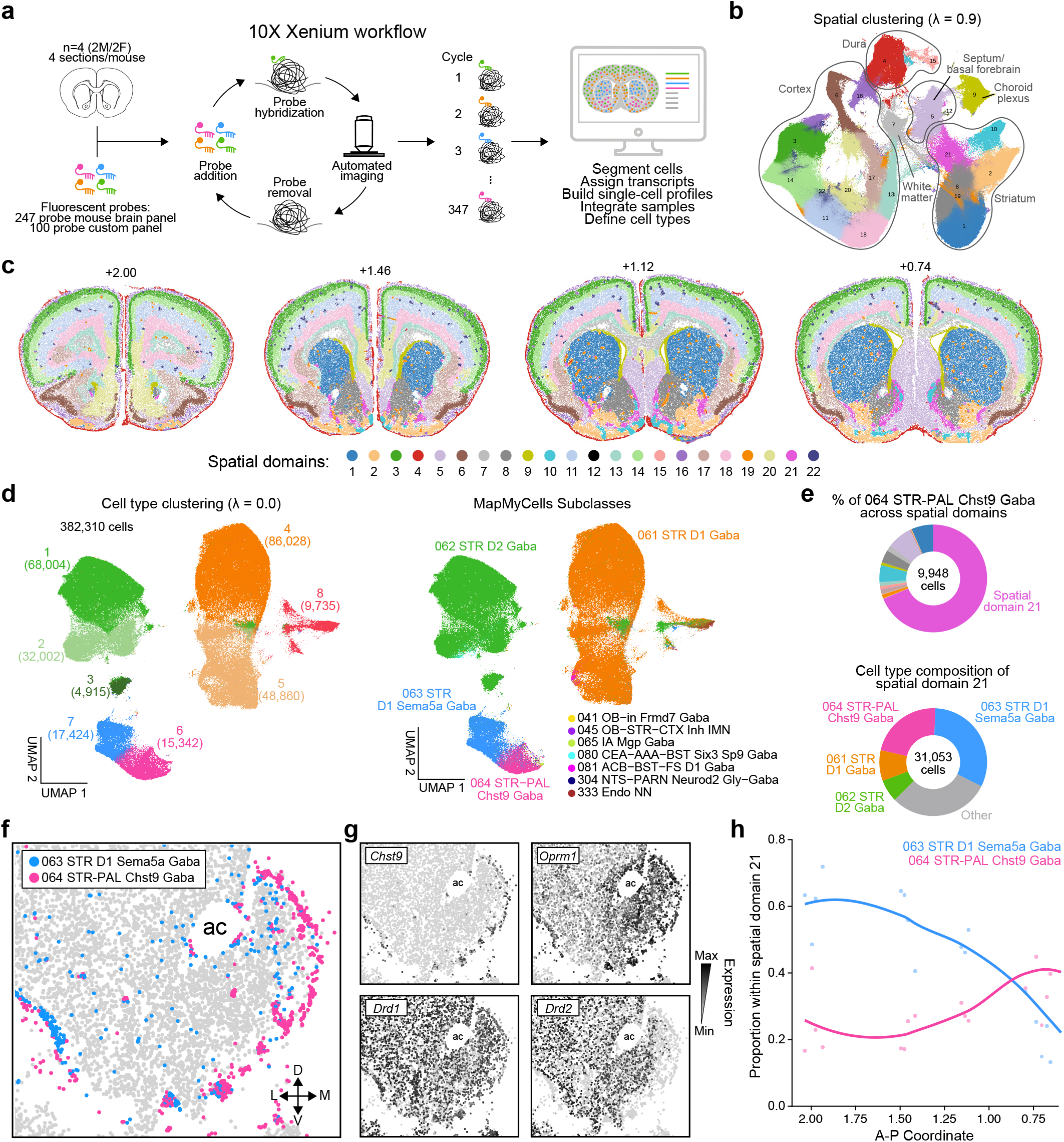
Chst9-MSNs define a spatially restricted µOR-enriched D1-MSN subpopulation at the nucleus accumbens shell border. **a**, Sixteen tissue sections from two male and two female mice (four sections per animal) were processed using the 10x Genomics Xenium spatial transcriptomics platform with a 347-gene panel. **b**, Spatial clustering (lambda = 0.9, number of neighbors (k) = 30, PCs = 30, and resolution = 1.0) of nuclei identified 22 spatial domains across the sampled tissue. **c**, Representative spatial plots showing the distribution of spatial domains across four sections from a single mouse. **d**, UMAP visualization of medium spiny neuron (MSN) clustering (lambda = 0.0, number of neighbors (k) = 30, PCs = 30, and resolution = 1.0) identified eight MSN cell types (left). MSN clusters were annotated using the MapMyCells tool from the Allen Institute (right). **e**, Quantification of Chst9-MSN localization across spatial domains revealed that spatial domain 21 contained approximately 70% of Chst9-MSNs (top). Quantification of MSN subtypes within spatial domain 21, demonstrating enrichment of Sema5a-MSNs and Chst9-MSNs (bottom). **f**, Spatial localization of Chst9-MSNs and Sema5a-MSNs in one representative tissue section. **g**, Spatial expression maps of *Chst9, Oprm1, Drd1*, and *Drd2*. **h**, The proportions of Chst9-MSNs and Sema5a-MSNs within spatial domain 21 from anterior to posterior sections.

Following quality control and sample integration, we retained 1,544,589 high-quality cells across 16 tissue sections. To identify spatial domains, we performed spatial clustering analysis with a high spatial location weight (λ = 0.9), which identified 22 molecularly distinct spatial domains (**Fig. 2b**, see methods for details on clustering parameters). The relative prevalence of identified domains varied across anterior to posterior positions (**Fig. 2c**). To define the cell type composition of these spatial domains, we performed transcriptional clustering without consideration of spatial location (λ = 0), which identified 39 cell types including several MSN populations (**Fig. S2a**). Using the Allen Institute’s MapMyCells tool, we identified the major subclasses of cell types as MSN, GABAergic, glutamatergic, glial, and vascular (**Fig. S2b**). Subclustering of the MSN subclasses identified eight MSN populations, which were then annotated using the MapMyCells tool (**Fig. 2d**). MapMyCells classified clusters 4 and 5 as D1-MSNs and clusters 1, 2, and 3 as D2-MSNs. Examination of their spatial distributions revealed that clusters 4 and 5 were enriched in the dorsal and ventral striatum, respectively, while clusters 1 and 2 exhibited similar dorsal and ventral distributions (**Fig. S2c**). In contrast, cluster 3 represented a sparse D2-MSN population localized to the NAc core and dorsal striatum. Clusters 6 and 7 corresponded to Chst9-MSNs and Sema5a-MSNs, respectively, and exhibited overlapping spatial distributions along the perimeter of the NAc shell.

We next assessed the distribution of Chst9-MSNs across the 22 spatial domains and found that this population is highly enriched in spatial domain 21, with nearly 70% of all Chst9-MSNs localizing to this domain (**Fig. 2e**). Additionally, spatial domain 21 was composed predominantly of Sema5a-MSNs and Chst9-MSNs. Consistent with previous reports localizing *Chst9* expression to the medial and ventral borders of the NAc shell^29^, spatial domain 21 is organized in discrete clusters and streaks along the NAc boundary at multiple anterior-posterior levels (**Fig. 2c**). Similarly, Chst9-MSNs and Sema5a-MSNs exhibit closely associated spatial distributions concentrated along the NAc shell border (**Fig. 2f**). Consistent with our snRNA-seq findings and prior studies, *Chst9* expression overlapped with *Oprm1* expression (**Fig. 2g**). In addition, Chst9-MSNs expressed *Drd1* and were localized to pockets along the NAc border that lack *Drd2* expression. Finally, to determine how the distribution of Chst9-MSNs varies along the anterior-posterior axis of the NAc, we assigned each tissue section a Bregma coordinate and quantified the proportion of Sema5a- and Chst9-MSNs across coordinates. This analysis revealed that Chst9-MSNs within spatial domain 21 were less abundant at anterior coordinates and became progressively enriched at more posterior levels (**Fig. 2h**).

### Chst9-MSNs predominantly project to the ventral pallidum

Although our snRNA-seq and spatial transcriptomic analyses indicate that Chst9-MSNs represent a subtype of *Drd1*-expressing MSNs distributed mostly along the perimeter of the NAc shell, their downstream projection targets have not been identified. To enable selective labeling and manipulation of Chst9-MSNs, we generated a knock-in rat line expressing Cre recombinase and tdTomato in frame with the *Chst9* gene (**Fig. 3a**). To define the anatomical targets of this population, we performed bilateral injections of AAV9-hSyn-DIO-eGFP in *Chst9*^*Cre-tdTomato*^ rats and analyzed GFP-labeled fibers by immunohistochemistry (**Fig. 3b**). GFP expression was localized to Chst9-MSN cell bodies in the NAc shell, confirming selective viral targeting of this population (**Fig. 3c**). Examination of sagittal sections co-labeled for substance P, a commonly used marker for VP boundaries^36^, revealed prominent GFP-positive projections to the VP (**Fig. 3d, e**). To further characterize the projection pattern, we examined coronal sections containing reported D1-MSN projection targets^37–42^, including the VP, lateral hypothalamus (LH), and VTA. This analysis confirmed robust GFP labeling in the VP (**Fig. 3e**). Sparse GFP-positive fibers were also observed in the LH (**Fig. S4a**), whereas no detectable GFP-positive projections were observed in the VTA (**Fig. S4b**). Together, these findings indicate that Chst9-MSNs exhibit a strong projection bias to the VP, which distinguishes this *Drd1*+ MSN subpopulation from canonical D1-MSNs that form the direct pathway out of the striatum.

**Figure 3.**
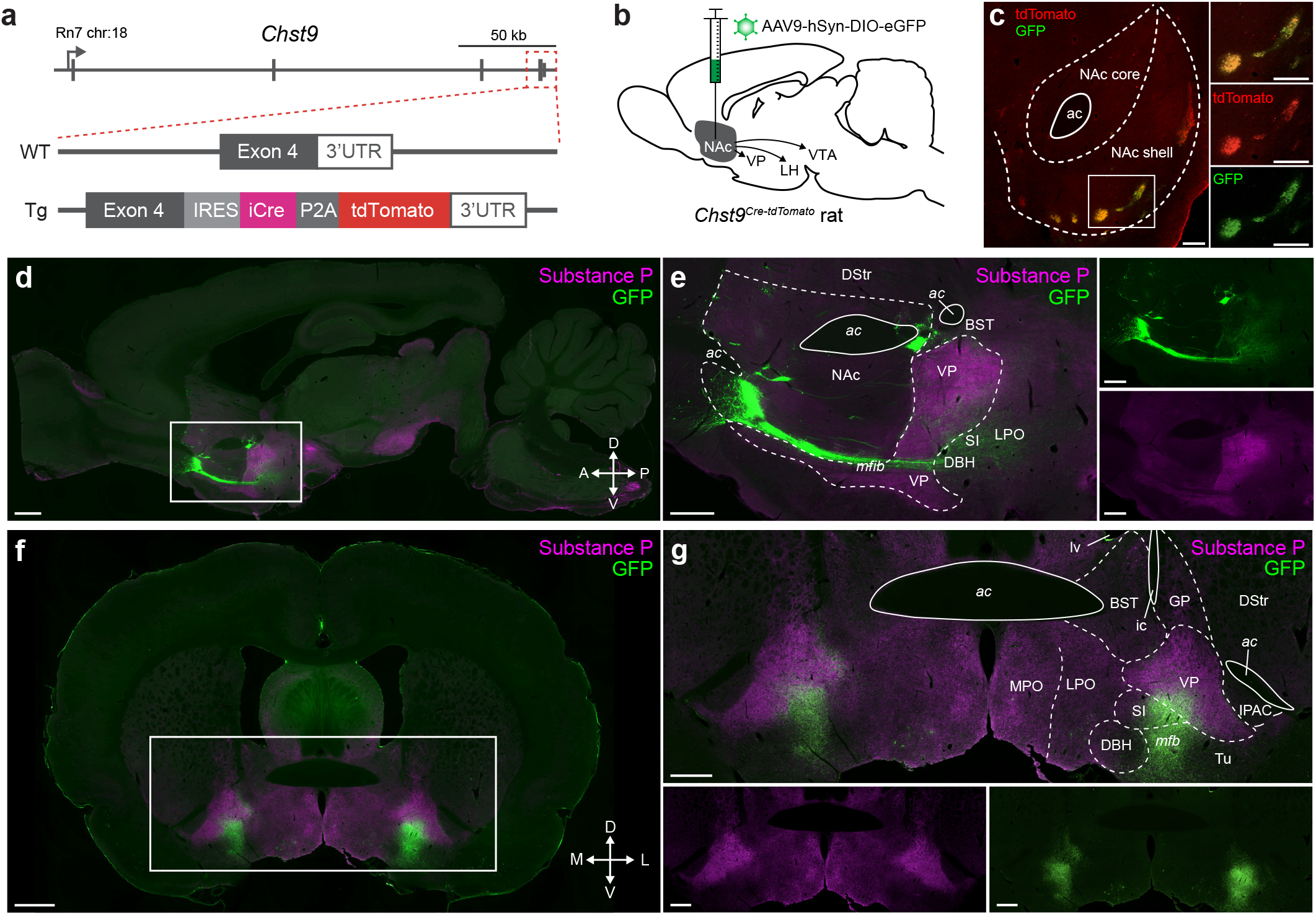
Projection mapping of Chst9-MSNs using a *Chst9*^*Cre-tdTomato*^ transgenic rat. **a**, Schematic of the *Chst9* locus showing insertion of the IRES-NLS-iCre-P2A-tdTomato cassette downstream of exon 4. **b**, Schematic of the approach for anterograde tracing of Chst9-MSN projections. **c**, Immunohistochemistry targeting tdTomato (red) and GFP (green) shows colocalization within NAc cell bodies; scale bar = 250 μm. **d**, Representative sagittal section showing GFP-labeled Chst9-MSN projections from the NAc to the ventral pallidum (VP); green = GFP, magenta = Substance P; scale bar = 1000 μm. **e**, Higher-magnification image of the region outlined in d showing GFP-positive fibers in the VP. DStr, dorsal striatum; ac, anterior commissure; BST, bed nucleus of the stria terminalis; VP, ventral pallidum; NAc, nucleus accumbens; mfb, medial forebrain bundle; SI, substantia innominata; LPO, lateral preoptic area; DBH, horizontal limb of the diagonal band; scale bar = 500 μm. **f**, Representative coronal section showing dense GFP-labeled terminals in the VP; scale bar = 1000 μm. **g**, Higher-magnification image of the region outlined in f. ac, anterior commissure; lv, lateral ventricle; BST, bed nucleus of the stria terminalis; ic, internal capsule; GP, globus pallidus; DStr, dorsal striatum; MPO, medial preoptic area; LPO, lateral preoptic area; SI, substantia innominata; DBH, horizontal limb of the diagonal band; VP, ventral pallidum; mfb, medial forebrain bundle; Tu, tubercle; IPAC, interstitial nucleus of the posterior limb of the anterior commissure.

### Chst9-MSNs are directly inhibited by opioids

Previous studies examining the effects of µOR activation on MSN electrophysiology have reported mixed findings, largely within broad D1- or D2-MSN populations^43–45^. For example, the µOR agonist DAMGO produces region- and cell type-specific effects in the striatum, including subtle increases in D1-MSN excitability in the NAc^46^. However, given the substantial molecular and functional heterogeneity within the D1-MSN class, these population-level effects may obscure responses of specific subtypes. Because Chst9-MSNs express high levels of *Oprm1*, we next examined the effects of µOR activation on their intrinsic electrophysiological properties. To this end, we performed whole-cell patch clamp recordings from identified tdTomato+ neurons in brain slices containing the NAc from *Chst9*^*Cre-tdTomato*^ rats (**Fig. 4a, b**). To isolate cell-autonomous effects of µOR activation, synaptic transmission was blocked by inclusion of D-APV (5µM), NBQX (10µM), picrotoxin (100µM), and SCH50911 (1µM) in the circulating artificial cerebrospinal fluid.

**Figure 4.**
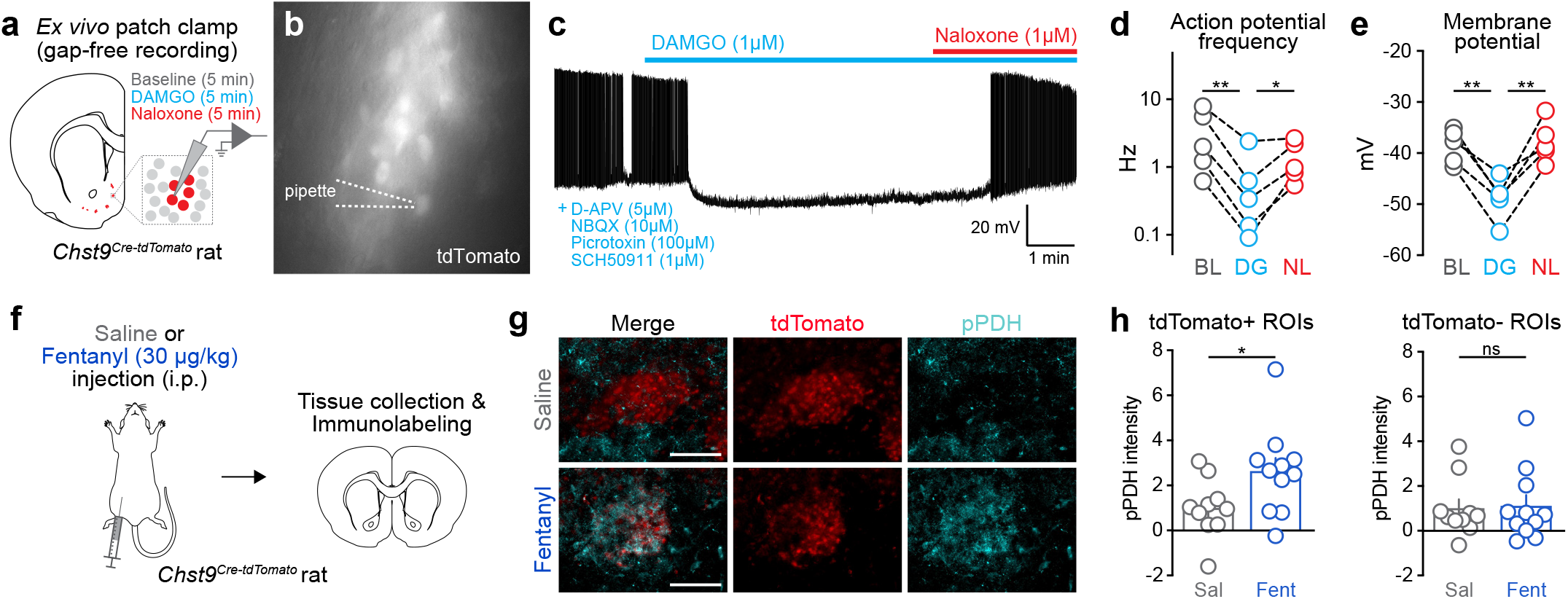
Chst9-MSNs are inhibited by µOR activation. **a**, Schematic of ex vivo slice electrophysiology recording approach in *Chst9*^*Cre-tdTomato*^ rats. **b**, Representative 40x image of a patched tdTomato+ (*Chst9*+) neuron visualized by tdTomato fluorescence. **c**, Representative trace demonstrating a decrease in action potential frequency and membrane potential following DAMGO (DG) application. Recordings were performed in the presence of synaptic blockers to isolate the contributions of Chst9-MSN µORs. Silenced activity was reversed by bath application of the µOR antagonist naloxone (NL). **d**, Significant decrease in action potential frequency following DAMGO is reversed by naloxone (Repeated measures ANOVA, F = 25.70, *p* = 0.0023, R^2^ = 0.86; Tukey’s multiple comparisons test). **e**, Significant decrease in membrane potential following DAMGO is reversed by naloxone (Repeated measures ANOVA, F = 26.25, *p* = 0.0006, R^2^ = 0.87; Tukey’s multiple comparisons test). N = 5 cells/rats. **f**, Schematic of in vivo experimental design. *Chst9*^*Cre-tdTomato*^ rats received intraperitoneal saline or fentanyl (30 µg/kg) and were perfused 1 h later for immunohistochemistry targeting phosphorylated pyruvate dehydrogenase (pPDH). **g**, Representative images showing tdTomato+ Chst9-MSNs (red) and pPDH immunoreactivity (cyan) following saline or fentanyl administration, scale bar = 100 µm. **h**, Acute fentanyl administration increased pPDH immunoreactivity in Chst9-MSNs (tdTomato+ ROIs) compared with saline-treated controls (unpaired t-test, t_19_ = 2.252, *p* = 0.0364). Fentanyl did not alter pPDH immunoreactivity in neighboring tdTomato-ROIs (unpaired t-test, t_19_ = 0.166, p = 0.8701). N = 10 saline-treated rats (5 male, 5 female) and 11 fentanyl-treated rats (5 male, 6 female). For visualization, pPDH immunoreactivity was normalized independently within tdTomato+ and tdTomato-ROIs to the mean of the corresponding saline group. \**p* < 0.05, \*\**p* < 0.01.

Current was injected in 10 pA increments to establish stable baseline firing, after which DAMGO (1µM) was bath-applied, followed by naloxone (1µM) to assess reversibility. DAMGO produced a robust reduction in action potential frequency that was reversed by naloxone, indicating a µOR-dependent effect (**Fig. 4c, d**). DAMGO also hyperpolarized the membrane potential, an effect that was similarly reversed following naloxone application (**Fig. 4e**). These findings demonstrate that µOR activation directly inhibits Chst9-MSN excitability. In contrast to the subtle excitatory effects previously reported in broad D1-MSN populations^46^, Chst9-MSNs exhibit a pronounced inhibitory response to µOR activation. These results provide functional evidence that the high levels of *Oprm1* expression observed in Chst9-MSNs translate into a robust physiological sensitivity to µOR activation.

To assess whether Chst9-MSNs are inhibited by opioids *in vivo*, rats received intraperitoneal (IP) injections of saline or fentanyl (30 µg/kg; **Fig. 4f**). In coronal brain sections containing the NAc, we performed immunohistochemistry for phosphorylated pyruvate dehydrogenase (pPDH; a recently described marker of neuronal inhibition^47^), to quantify neuronal silencing by fentanyl in Chst9-MSNs. Relative to saline-treated controls, acute fentanyl administration increased pPDH immunoreactivity in Chst9-MSNs (**Fig. 4g, h**), consistent with opioid-induced inhibition of this population. Importantly, no significant change in pPDH immunoreactivity was detected in neighboring tdTomato– cells. Together, these findings show that Chst9-MSNs are selectively inhibited by both direct µOR activation ex vivo and systemic fentanyl administration in vivo.

#### µOR signaling in Chst9-MSNs is required for opioid reward learning

Given that Chst9-MSNs disproportionately express µORs and are directly sensitive to opioids, we next addressed whether opioid signaling in these neurons is required for opioid reward learning. To selectively disrupt *Oprm1* expression in Chst9-MSNs, we generated a Cre-dependent AAV9 vector encoding SaCas9 and a single guide RNA (sgRNA) targeting exon 2 of *Oprm1*, which is retained across canonical µOR isoforms (**Fig. 5a, b, Fig. S5**)^48^. An identical virus encoding an sgRNA targeting the bacterial gene *lacZ* was used as a control. Viruses were bilaterally injected into the NAc shell of female *Chst9*^*Cre-tdTomato*^ rats. Immunohistochemistry for µOR and a virally encoded hemagglutinin (HA) tag revealed a greater than 60% reduction in µOR intensity in HA+ cell clusters in *Oprm1*-targeted rats relative to *lacZ* controls (**Fig. 5d, e**; unpaired t-test, *p* < 0.0001), confirming effective µOR depletion in virally transduced cells.

**Figure 5.**
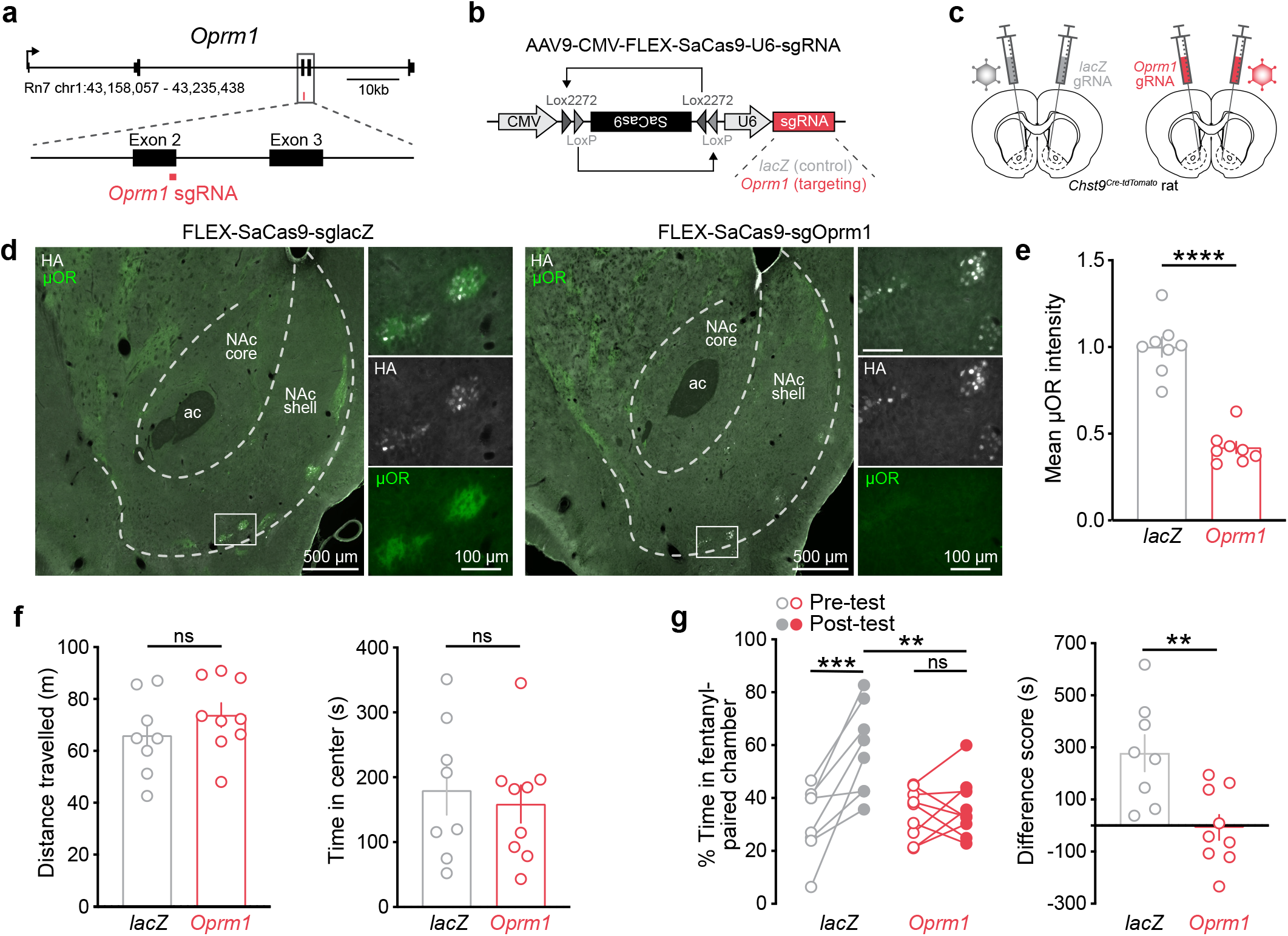
Cre-dependent µOR knockout in Chst9-MSNs prevents fentanyl reward learning. **a**, *Oprm1* locus showing single guide RNA (sgRNA) target site in Exon 2. **b**, Cre-dependent SaCas9 viral construct used for *Oprm1* knockout. **c**, Targeting strategy for viral delivery to the NAc shell in *Chst9*^*Cre-tdTomato*^ rats. **d**, Representative images of µOR protein (green) and HA (white) in control (*lacZ*, left) and *Oprm1* knockout (right) animals. **e**, Quantification of µOR intensity in HA+ cells in the NAc shell validates successful knockout of *Oprm1* (unpaired t-test, t_15_ = 8.791, *p* < 0.0001). **f**, Open field test. *Oprm1* knockout did not affect total distance travelled (left; unpaired t-test, t_15_ = 1.088, *p* = 0.2940) or time in center of field (right; unpaired t-test, t_15_ = 0.4393, *p* = 0.6667). **g**, Knock-out of *Oprm1* in *Chst9*+ cells abolishes fentanyl place preference. Left, percent time spent in the fentanyl-paired chamber during pre-test and post-test sessions (Two-way repeated-measures ANOVA, Šídák multiple comparisons test). A significant effect of time (F_(1,15)_ = 18.32, *p* = 0.0007, ηp^2^ = 0.55) and a significant sgRNA x time interaction were observed (F _(1,15)_ = 13.43, *p* = 0.0023, ηp^2^ = 0.47). *lacZ* animals exhibited significant fentanyl CPP following conditioning (*p* = 0.0003), whereas *Oprm1* knockout animals did not (*p* = 0.9866). Post-conditioning time spent in the fentanyl-paired chamber was significantly lower in *Oprm1* knockout animals relative to controls (*p* = 0.0063). Right, Difference scores (post-test minus pre-test time in the fentanyl-paired chamber) were lower in *Oprm1* knockout animals compared to *lacZ* controls (t_15_ = 3.410, *p* = 0.0039). \*\**p* < 0.01, \*\*\**p* < 0.001

We next performed fentanyl (15 µg/kg) conditioned place preference (CPP) to determine whether *Oprm1* deletion in Chst9-MSNs alters opioid reward learning. Preference for the fentanyl-paired chamber was quantified across pre- and post-conditioning sessions. Baseline locomotor activity did not differ between groups prior to conditioning (**Fig. 5f**). For fentanyl CPP, group comparisons revealed a significant interaction between sgRNA treatment and conditioning session. Post hoc analyses showed that *lacZ* control animals exhibited increased preference for the fentanyl-paired chamber following conditioning (*p* = 0.0003), whereas *Oprm1*-targeted animals failed to develop CPP (*p* = 0.9866, **Fig. 5g**). Additionally, time spent in the fentanyl-paired chamber following conditioning was significantly lower in *Oprm1* knockout animals relative to controls (*p* = 0.0063). Consistent with this effect, CPP scores (% time in fentanyl-paired chamber during post-test minus pre-test) were significantly lower in *Oprm1*-targeted animals compared to *lacZ* controls (*p* = 0.0039, **Fig. 5g**). Together, these findings demonstrate that µOR signaling in Chst9-MSNs is required for fentanyl reward learning.

## DISCUSSION

Recent large-scale sequencing studies have expanded our understanding of cellular diversity within the NAc, revealing multiple transcriptionally distinct MSN subtypes across species^20–26,31,32^. However, the functional relevance of many genetically defined populations remains poorly understood. Here, we identify Chst9-MSNs as a transcriptionally, spatially, and functionally distinct D1-MSN subpopulation enriched for µOR expression. Although relatively sparse, Chst9-MSNs account for a substantial proportion of accumbal *Oprm1* expression, form discrete clusters along the NAc shell border, and exhibit a conserved molecular identity across species. Unlike the broader projection patterns described for NAc shell D1-MSNs, Chst9-MSNs project predominantly to the VP, suggesting specialized circuit organization. Functionally, Chst9-MSNs are strongly inhibited by µOR activation, and selective deletion of *Oprm1* from this population abolishes fentanyl CPP, demonstrating that µOR signaling in this discrete MSN subtype is required for opioid reward learning.

Together, these findings support growing evidence that the classical D1/D2 framework incompletely captures the cellular and functional diversity of the NAc^20,22,24,49–55^. Although MSNs have traditionally been divided into *Drd1*- and *Drd2*-expressing populations associated with broadly opposing behavioral functions, manipulations of these populations often produce inconsistent behavioral outcomes, likely reflecting substantial heterogeneity within each class^56–61^. Existing models similarly treat D1-MSNs as a relatively uniform substrate for opioid reward. Previous studies of D1-MSNs have shown that µOR activation produces net excitatory effects in whole-cell recordings^46^ and an increase in neuronal activation markers^62^. In contrast, Chst9-MSNs are robustly inhibited by µOR activation despite their D1 identity, suggesting that opioid reward may involve suppression of a specialized D1-MSN subtype rather than uniform engagement of D1 circuitry. Although recent transcriptomic and spatial studies have identified sparse “eccentric”^31^ and “atypical”^20^ MSN populations with molecular and anatomical features resembling Chst9-MSNs^24,32^, these studies remained largely descriptive. Here, we directly link transcriptional identity to molecular, anatomical, physiological, and behavioral specialization, suggesting that behavioral functions attributed to broad D1-MSN populations may arise from discrete genetically defined cell types.

Previous studies established that µOR signaling in the NAc is a critical regulator of reward-related behaviors^8,10,63–72^. Direct infusion of opioid agonists into the NAc is reinforcing in self-administration assays^8–10,12^, whereas disruption of µOR signaling through global *Oprm1* deletion^1^ or selective lesioning of *Oprm1*-expressing NAc neurons^14^ reduces opioid reward and seeking behaviors. However, the cellular substrates mediating these effects remained unclear. Recent studies have begun to implicate transcriptionally defined MSN populations in opioid-related behaviors^34^, and our findings extend this work by identifying Chst9-MSNs as a sparse but µOR-enriched population that exhibits robust µOR-mediated inhibition and is required for fentanyl reward learning. These findings identify Chst9-MSNs as a major cellular substrate through which accumbal µOR signaling contributes to opioid reward, and suggest that this population may have been the key functional target in previous NAc-wide µOR manipulations.

The strong VP projection bias of Chst9-MSNs provides a potential circuit mechanism through which µOR signaling influences opioid reward. As GABAergic projection neurons that are robustly inhibited by µOR activation, opioid exposure may reduce Chst9-MSN-mediated inhibition of VP targets. This model is consistent with recent work showing that inhibition of VP-projecting D1-MSNs is rewarding and promotes dopamine release^40^. Chst9-MSNs share several features with this population, including predominant projection to the VP and relatively low *Drd1* expression, raising the possibility that µOR-mediated suppression of Chst9-MSNs engages similar circuitry. This interpretation is further supported by the established role of the VP as a critical regulator of reward and motivation^53,73,74^. In addition to serving as a major output target of the NAc, the VP contains an opioid “hedonic hotspot” in its posterior region capable of enhancing reward processing^74^. Additionally, the VP projects to mesolimbic dopamine circuits and contributes to natural and opioid reward processing, relapse to opioid seeking, and locomotor sensitization to opioids^52,75–79^. Thus, despite comprising only a small fraction of accumbal neurons, Chst9-MSNs may exert disproportionate influence over opioid reward through their enrichment for *Oprm1* and connectivity with VP circuitry.

These findings may also help resolve a longstanding paradox in striatal organization, which is that the classic association of D1-MSNs with the direct pathway and D2-MSNs with the indirect pathway^15^ does not strictly apply in the NAc^80^. Because Chst9-MSNs express *Drd1*, they are likely included in Cre-dependent approaches targeting the broader D1-MSN population. Importantly, recent developmental lineage studies show that *Chst9*+ neurons arise from the same immature GABAergic precursor population that gives rise to all striatal MSN subclasses, first emerging during the early postnatal migration of lateral ganglionic eminence-derived neurons^81^. This observation supports an emerging cross-species consensus that striatal cell type organization is much more complex than previously anticipated, with numerous subpopulations of both D1- and D2-MSNs that possess distinct spatial, molecular, and circuit properties^20–23,25,26,29,82^.

The translational relevance of Chst9-MSNs is supported by their conservation across rodents, non-human primates, and humans, where homologous cell types consistently exhibit robust *Oprm1* expression. This conservation suggests that the circuit and behavioral functions identified here may extend beyond rodents and could contribute to human opioid use disorder. Human genetic studies have implicated *OPRM1* in opioid-related phenotypes, including opioid use disorder and opioid sensitivity^83–86^, and our findings suggest that these associations may reflect effects driven by specialized *OPRM1*-enriched populations rather than across all accumbal neurons. Notably, several other genes enriched in Chst9-MSNs, including *TSHZ1*^*87*^, *GRM8*^*88,89*^, *FOXP2*^*90,91*^, and *CHST9* itself^92,93^ have also been linked by GWAS to neuropsychiatric and substance use-related phenotypes, suggesting that this cell type may represent a convergence point for genetic risk relevant to addiction. More broadly, these findings demonstrate how integrating single cell transcriptomics with circuit, physiological, and behavioral approaches can connect human genetic discoveries to specific neural cell types. Together, our results establish Chst9-MSNs as a conserved cellular substrate for opioid reward and provide a revised framework for understanding how genetically defined neuronal populations contribute to reward-related behaviors.

## METHODS

### Animals

All animal experiments were performed in accordance with the University of Alabama at Birmingham Institutional Animal Care and Use Committee (animal protocol no. 23174). Sprague-Dawley adult male and female rats (90 days old, snRNA-seq) were purchased from Charles River Laboratories (Wilmington, MA, USA). Adult C57BL/6J mice (11-12 weeks old) were purchased from Jackson Labs and used for Xenium Spatial Transcriptomics. The *Chst9*^*Cre-tdTomato*^ rat line on a Sprague-Dawley background was used for all viral targeting, electrophysiology, histological, and behavioral experiments. Rats were co-housed in pairs in plastic-filtered cages with wooden chewing block enrichment in an American Association for Accreditation of Laboratory Animal Care (AAALAC)-approved animal care facility maintained between 22° and 24°C on a 12-hour light/ dark cycle with ad libitum access to food (Lab Diet SL3Z Irradiated rat chow) and water. Bedding and enrichment were changed weekly by animal resources program staff. Animals were randomly assigned to experimental groups. All animals were handled by investigators for 5 to 7 days before behavioral testing.

### Single-nucleus RNA sequencing

#### Tissue collection

Rats were euthanized by live decapitation and whole brains were rapidly extracted and briefly flash frozen in 2-methylbutane on dry ice and stored at -80°C until further processing. 2mm-deep bilateral NAc tissue punches were collected at approximately Bregma +2.42 in a cryostat at -18°C. Punches were immediately placed in pre-chilled sample dissociation tubes (10x Genomics, PN-2000564) and kept at -80°C until nuclei isolation.

#### Single-nuclei dissociation

Nuclei were isolated using the 10x Genomics Nuclei Isolation Kit (PN-1000493) according to manufacturer’s instructions, with all steps performed on ice. Tissue was mechanically dissociated in lysis buffer, filtered through a nuclei isolation column, and underwent centrifugation and debris removal steps. Purified nuclei were washed and resuspended in 1x PBS containing 1% BSA (Thermo Scientific #37525) and 2.5% RNAse Inhibitor (10x Genomics, 2000565), and then kept on ice. Nuclei concentration and viability were quantified via 7-aminoactinomycin D (7-AAD) staining and observed with a Nikon TiS inverted fluorescence microscope.

#### Library preparation, GEM generation, and single-nuclei RNA sequencing

Single-nucleus libraries were generated using the Chromium GEM-X Single Cell 3’ Kit v4 according to manufacturer’s instructions (10x Genomics, PN-1000691). Following isolation, nuclei were concentrated to 1,000 nuclei per μL. A total of 8,000 nuclei pooled equally from one male and one female sample were loaded onto individual wells of a GEM-X 3’ Chip (PN-2001097) for GEM generation and barcoding using the Chromium Controller (PN-120270). cDNA amplification and library construction were performed using the Single Cell 3’ GEM Kit v4 (PN-1000693), the Library Construction Kit C (PN-1000694), and SPRIselect bead-based cleanup (Beckman Coulter, B23318) according to manufacturer’s protocol. The cDNA and library quality were assessed using a Bioanalyzer High Sensitivity DNA assay (Agilent Technologies, Santa Clara, CA, USA). A total of 64,260 nuclei were captured across 6 GEM (gel bead in emulsion) wells in total using the 10x Chromium X. Each GEM well contained nuclei from 1 male and 1 female for a total of 6 rats per sex (mean = 10,710 nuclei per GEM well). Libraries were sequenced on an Illumina NovaSeq 6000 at the UAB Heflin Genomics Core to an average depth of 232,782,011 reads per library, which equated to an average depth of 22,762 reads per nucleus.

#### snRNA-seq analysis

Raw sequencing data for each GEM well were processed and aligned to the rat genome (mRatBn7.2) in Cell Ranger^94^ version 9.0.1 on the Cheaha high-performance computing cluster at the University of Alabama at Birmingham, with the associated Ensembl gene transfer format file (version 113) modified to add annotation for the *Xist* gene. Filtered Cell Ranger outputs were analyzed with Seurat^95^ version 5.2.1 in R^96^ version 4.4.0. Ambient RNA contamination was estimated and removed using SoupX^97^ version 1.6.2. Cells with fewer than 200 genes or greater than 2% mitochondrial transcripts were excluded from downstream analysis. For each GEM well, counts were normalized using Seurat’s NormalizeData function with a scale factor of 10,000, and log transformed. Heterotypic doublets were identified and removed using scDblFinder^98^ version 1.18.0 with the default expected doublet rate of 1% per 1,000 recovered cells. To infer the sex of cells from pooled GEM wells, *Xist* expression was used as a binary classifier, with cells expressing *Xist* (> 0 counts) classified as female and cells lacking detectable *Xist* expression (= 0) classified as male.

Samples were integrated using Harmony^99^ version 1.2.3 based on principal component analysis (PCA) embeddings computed from the top 50 principal components (PCs). Downstream analyses were performed on the integrated dataset. Gene expression values were scaled to a mean of 0 and variance of 1 across cells using Seurat’s ScaleData function. Dimensionality reduction was performed using PCA retaining 21 PCs, followed by UMAP construction using the same 21 PCs. Cells were clustered under a graph-based approach by identifying k-nearest neighbors in PCA space and applying the Louvain algorithm with a resolution of 0.5. These parameters were chosen by iterating through combinations of 17-23 PCs and resolution of 0.2-0.6 and selecting the parameter set that best recapitulated known NAc cell populations.

To refine cell class annotation, clusters were mapped using the Allen Institute’s Whole Mouse Brain taxonomy using MapMyCells web interface (RRID:SCR_024672). After removal of low-quality or ambiguous populations, dimensionality reduction and clustering were repeated using the same parameters. Cells were considered low quality or ambiguous if they belonged to clusters mapping inconsistently across multiple MapMyCells classes or subclasses with low mean bootstrapping probabilities and had low unique molecular identifier (UMI) and gene counts. Additionally, clusters corresponding to subclasses 057 NDB-SI-MA-STRv (within class 08 CNU-MGE GABA) and 085 SI-MPO-LPO Lhx8 Gaba (within class 11 CNU-HYa GABA) were excluded due to their anatomical localization outside the NAc. DEGs between MSN populations were identified using Seurat’s FindAllMarkers function (Wilcoxon rank-sum test) with Bonferroni correction for multiple comparisons. Genes were considered significant if they had an adjusted p value < 0.05, average log_2_ fold change > 0.5, and were detected in >50% of cells within that cluster.

### Cross-species cell-type mapping

To evaluate the conservation of NAc MSN populations across species, we performed cross-species cell-type correspondence analysis using Allen Institute’s MapMyCells^28^, which allows the use of reference cell-type taxonomies to identify transcriptionally corresponding populations across independently generated snRNA-seq datasets. The MapMyCells built-in mouse reference taxonomy^30^, as well as published reference datasets from NHP striatum^24^ and human striatum^26^ were used for cross-species comparison. NHP and human datasets were downloaded as processed datasets, annotated according to published cell-type classifications, and restricted to MSN populations prior to cross-species mapping. The rat NAc snRNA-seq dataset from Figure 1 was subset to six MSN populations (Drd1-MSN-1, Drd1-MSN-2, Drd2-MSN-1, Drd2-MSN-2, Chst9-MSN, and Sema5a-MSN).

To enable cross-species comparison, rat genes were converted to orthologous mouse, NHP, and human gene identifiers using a one-to-one ortholog table. After genes lacking unique orthologs across species were excluded, a total of 13,859 genes remained for mapping. Count matrices containing conserved orthologous genes were converted to AnnData format for compatibility with MapMyCells. Cross-species mapping was performed using the hierarchical correlation mapping algorithm in the MapMyCells web interface (mouse) or standalone cell-type mapper tool (NHP, human) with raw count normalization and, for the NHP and human custom taxonomies, on-the-fly marker gene selection. Rat MSN populations were independently mapped to NHP and human MSN reference taxonomies. Mapping confidence was assessed using bootstrap probabilities, and cells with low-confidence assignments (bootstrap assignment probability < 0.5) for mouse class or subclass were excluded from downstream visualization for clarity. We also removed mouse classes or subclasses with < 100 mapped cells. The resulting cell-type annotations were integrated into the rat Seurat object for downstream analyses. To visualize correspondence between species, cell-type assignments were summarized by population abundance and displayed using alluvial plots generated with the R package ggalluvial version 0.12.5^100^. Complete analysis scripts and computational workflows are available on GitHub (see Data Availability).

### Xenium spatial transcriptomics

#### Sample collection

Two female and two male mice (11 weeks old) were used for spatial transcriptomics with the 10x Genomics Xenium platform. Mice were euthanized by live decapitation and whole brains were rapidly extracted and immediately flash frozen in 2-methyl butane on dry ice. Brains were kept at -80ºC until further processing. Brains were sectioned at 10μm on a Leica CM 1860 cryostat (Leica Biosystems, Deer Park, IL, USA). Four sections from each mouse spanning Bregma +2.04 mm and +0.60 mm were collected and placed on Xenium slides (10x genomics, PN-1000460). Slides were then stored at -80ºC for a maximum of three days before proceeding with the Xenium protocol.

#### In situ spatial transcriptomic profiling

The Xenium pipeline was conducted by the Single Cell and Flow Cytometry Core at the University of Alabama at Birmingham. Profiling was performed using the mouse gene expression panel (10x Genomics, PN-1000462) in addition to a custom gene panel (**Table S1**) for a total of 347 probes. The sections underwent fixation and permeabilization (protocol CG000579 Rev F) followed by probe hybridization, ligation, and amplification (protocol CG000749 Rev B). The slides were then placed in the Xenium analyzer where iterative rounds of fluorescent probe hybridization, imaging, and fluorophore cleavage were performed according to manufacturer’s protocol (protocol CG000584 Rev G). The instrument quantified and registered four-channel signal combinations to construct positional barcode matrices for subsequent quantification and computational analyses. In addition, multimodal segmentation was performed, including by boundary stain (ATP1A1, CD45, E-Cadherin), interior stain (18S rRNA), and/or nucleus expansion of a maximum of 5.0µm.

#### Data processing, spatial domain definition, and cell type annotation

Data processing was performed by the University of Alabama at Birmingham Biological Data Science Core using their Nextflow nf_xpatial pipeline using Nextflow version 24.04.3^101^. Briefly, nf_xpatial performed comprehensive preprocessing and quality assessment, including identification and removal of low-quality cells (where cells with fewer than 10 transcripts and fewer than 10 unique features were excluded), as well as quality-control evaluation pre and post filtering. The processed data was then area-normalized^102^, and integrated^99^ (Harmony, version 1.2.4) across samples to support comparative downstream analyses. Next, cell-level clustering and spatial domain clustering was performed with Seurat^95^ version 5.4.0 (single-cell clustering) and BANKSY^103^ version 1.5.10 (single-cell and spatial clustering). The Seurat based clustering was evaluated across the following parameters: 25 and 30 PCs and resolutions of 0.4-0.7. The BANKSY clustering was evaluated across the following parameters: use_agf = True (azimuthal Gabor filter), lambda: 0.0, 0.8, 0.9; k_geom: 30; PCs: 20 and 30; and resolutions ranging from 0.5 to 1.5. All clustering results were visualized with canonical markers and UMAPs.

For spatial domain clustering, based on interpretability of differentially expressed genes (DEGs) across spatial domains, we selected lambda = 0.9, number of neighbors (k) = 30, PCs = 30, and resolution = 1.0. This resulted in the identification of 22 spatial domains, which were consistently identified across samples, with the exception of spatial domain 12, which was detected only in the most posterior section from one mouse and corresponded to the preoptic area. For cell type clustering, lambda (weight given to spatial proximity) was reduced to 0 while other parameters were kept the same. After clustering, cell types were assigned using Allen Institute’s MapMyCells tool^28^. From the 39 cell type clusters, we subclustered the MSN populations (061 STR D1 Gaba, 062 STR D2 Gaba, 063 Sema5a, 064 Chst9, 061 D1, 060 D3 Folh1, 061 D1, 062 D2, 333 Endo). We repeated clustering with the same parameters and then removed 060 OT D3 Folh1 Gaba, 041 OB−in Frmd7 Gaba, 042 OB−out Frmd7 Gaba, 044 OB Dopa−Gaba, 040 OB Trdn Gaba, 045 OB−STR−CTX Inh IMN, and 039 OB Meis2 Thsd7b Gaba due to their anatomical locations in the Islands of Calleja and olfactory bulb. Subclustering of MSN subclasses identified eight MSN populations, which were then annotated using the MapMyCells tool. Clusters 4 and 5 represented dorsal and ventral D1-MSNs, respectively, while 1 and 2 were identified as dorsal and ventral D2-MSNs. Cluster 3 represented a sparse D2-MSN population localized to the NAc core and dorsal striatum. Clusters 6 and 7 corresponded to Chst9-MSNs and Sema5a-MSNs, respectively. Cluster 8 displayed a diffuse distribution throughout the striatum and mapped to multiple MSN subclasses as well as an endovascular population (333 Endo), suggesting that it likely represents segmentation-related artifacts rather than a distinct biological population.

### Generation and genotyping of *Chst9*^*Cre-tdTomato*^ knock-in rat line

_The Sprague-Dawley-*Chst9*_*IRES-iCre-P2A-tdTomato* (*Chst9*^*Cre-tdTomato*^) rat line was generated by Biocytogen (Waltham, MA, USA) using CRISPR/Cas9-mediated homology-directed repair in Sprague Dawley embryos (**Fig. S3**). An IRES-iCre-P2A-tdTomato cassette flanked by homology arms was inserted into the endogenous *Chst9* locus between exon 4 and the 3′UTR. Founder animals were screened by PCR and sequencing to confirm correct targeting and bred to establish the line. Upon receipt of the line at the University of Alabama at Birmingham, genotype was independently verified in-house by PCR amplification of the targeted locus and agarose gel electrophoresis.

#### Genomic DNA extraction from ear punch samples

Genomic DNA was extracted from ear punch samples using a proteinase K digestion and isopropanol precipitation method. Briefly, samples were incubated overnight in 55ºC with agitation in 500 µL digestion buffer containing Tris-HCl (pH 8.0), EDTA, NaCl, SDS, and Proteinase K (10 mg/ mL). Following digestion, samples were centrifuged to remove debris, and the supernatant was transferred to a fresh tube. DNA was precipitated by addition of an equal volume of isopropanol, followed by centrifugation and washing with 75% ethanol. DNA pellets were air-dried briefly and resuspended in distilled water. Samples were incubated at 55ºC for 2 h to facilitate DNA dissolution, and DNA concentration was determined using a NanoDrop One Microvolume UV-Vis Spectrophotometer (Thermo Scientific, Waltham, MA, USA) prior to downstream genotyping.

#### PCR Genotyping

Genomic DNA was amplified by PCR to distinguish wild-type and transgenic alleles of the *Chst9-IRES-iCre-P2A-tdTomato* locus. Separate PCR reactions were performed to detect the wild-type allele and the transgenic allele using allele-specific primer combinations. The wild-type allele was amplified using a forward primer targeting exon 4 of the *Chst9* gene (5’-AGAGGGTCAGCAAACTGTGTTATC-3’) and reverse primer targeting the 3’ untranslated region (5’-GCTTCCTTGGTGAATGTAACAGCTG-3’). The transgenic allele was amplified using the same *Chst9* exon 4 forward primer and a transgenic reverse primer targeting the IRES region of the inserted transgene (5’-ACATTGCCAAAAGACGGCAATATG-3’). PCR reactions were prepared using Q5 Hot Start High-Fidelity 2x Master Mix (New England Biolabs, M0492S), 10 µM forward and reverse primers, and genomic DNA template. Reactions were performed on a C1000 Thermal Cycler (Bio Rad, Hercules, CA, USA) using an initial denaturation at 98ºC for 30s, followed by 32 cycles of PCR (10 seconds at 98ºC for denaturation, 30 seconds at 66ºC for annealing, and 4 minutes at 72ºC for extension), with a final extension at 72ºC for 5 minutes. WT-F and WT-R primers result in a 587 bp product for a wild-type locus and a 3732 bp product for a transgenic locus; WT-F and Tg-R primers result in a single 393 bp PCR product for a transgenic locus. PCR products were separated by agarose gel electrophoresis for genotype determination. Wild-type amplicons were resolved on 1% agarose gel containing GelRed (Biotium, 41003) in TAE buffer, and transgenic-targeted PCR products were resolved on a 2% agarose gel to distinguish expected amplicon size. Samples were mixed with loading dye and run alongside an appropriate DNA ladder at 90 V. Amplicons were visualized using an Azure 600 Gel Imager (Azure Biosystems, Dublin, CA, USA).

### smRNA FISH

#### Tissue collection

*Chst9*^*Cre-tdTomato*^ rats were euthanized by live decapitation and whole brains were rapidly extracted and immediately flash frozen in 2-methyl butane on dry ice. Brains were kept at -80ºC until further processing. Brains were sectioned at 10 μm on a Leica CM 1860 cryostat (Deer Park, IL, USA) and sections were immediately placed onto Superfrost Plus Microscope Slides (Thermo Fisher Scientific, Waltham MA, USA). Slides were kept at -80ºC until proceeding with the smRNA FISH protocol.

#### smRNA FISH and image acquisition

smRNA FISH probing for *Chst9* (1149681-C4), *iCre* (423321-C2), and *tdTomato* (317041-C3) was performed according to manufacturer’s protocol (no. 323136 ACD Bio, Newark, CA, USA) for fresh frozen tissue. Tissue was submerged in ice-cold 4% PFA for 15 min, serially dehydrated with ethanol, and then treated with hydrogen peroxide followed by protease IV digestion. Slides were incubated with a combination of 2 probes (*Chst9* and *iCre* or *Chst9* and *tdTomato*). Probes were fluorescently labeled with Opal Dyes diluted 1:750 (Akoya Biosciences, Marlborough, MA, USA; Opal *570* assigned to *tdTomato* and *iCre* and Opal 690 assigned to *Chst9*). Sections were stained with 4′,6-diamidino-2-phenylindole (DAPI) and cover slipped with ProLong Glass Antifade Mountant (Thermo Fisher Scientific, P36984, Waltham MA, USA). All images were acquired on a Keyence-BZ800 microscope. Within an experiment, all images were acquired with the same acquisition settings for a given objective and channel. 20x-magnification images of one striatal hemisphere were obtained across eight animals. Images were stitched using Keyence Image Analyzer software.

#### smRNA FISH image analysis

For puncta analysis to determine colocalization of *Chst9* and *tdTomato* or *iCre*, thresholds were defined in QuPath v0.3.2^104^ and held consistent for each channel across animals. In downstream analyses, K-means clustering was used to identify relevant threshold values for puncta counts. R^96^ version 4.4.0 was used to quantify overlap of puncta within a given nuclear ROI.

### Plasmid construction and preparation of SaCas9-sgRNA Vectors

#### sgRNA Design and Cloning

A single guide RNA (sgRNA) targeting exon 2 of *Oprm1* was designed using CHOPCHOP^105^ (RRID:SCR_015723) and evaluated for predicted specificity using NCBI BLAST (RRID:SCR_004870). An sgRNA targeting the bacterial *lacZ* gene served as a non-targeting control. *Oprm1* sgRNA sequence: 5’-GTGGCAACCACAAAATACAGG-3’; *lacZ* sgRNA sequence: 5’-GTATGCTTCGCTGCCAACGT-3’. Complementary sense and antisense oligonucleotides containing BsaI-compatible overhangs were synthesized (Sigma-Aldrich), annealed, phosphorylated using T4 Polynucleotide Kinase (New England Biolabs, M0201S), and ligated into BsaI-digested SaCas9 expression vectors using T4 DNA Ligase (New England Biolabs, M0202S). sgRNAs were cloned into both the constitutive vector pAAV-CMV::NLS-SaCas9-NLS-3xHA-bGHpA;U6::BsaI-sgRNA (Addgene #61591)^106^ and the Cre-dependent vector pAAV-FLEX-SaCas9-U6-sgRNA (Addgene #124844)^107^ using this strategy.

#### Colony Screening and Plasmid Verification

Ligation products were transformed into Stbl3 competent *E. coli* cells and plated on selective agar. Positive colonies were identified by colony PCR using a U6 forward primer (5’-TTTCTTGGGTAGTTTGCAGTTTT-3’) and corresponding sgRNA antisense oligonucleotide, with amplification performed using OneTaq 2x Master Mix (New England Biolabs, M0482S) on a C1000 Thermal Cycler (Bio Rad, Hercules, CA, USA) with the following thermocycler program: initial denaturation for 30 seconds at 94ºC, 32 cycles of PCR (15 seconds at 94ºC for denaturation, 20 seconds at 59ºC for annealing, and 15 seconds at 68ºC for extension), and final extension of 5 minutes at 72ºC. PCR products were resolved on a 2% agarose gel containing GelRed (Biotium, 41003) in TAE buffer. Samples were mixed with loading dye and run alongside an appropriate DNA ladder at 90 V. Amplicons were visualized using an Azure 600 Gel Imager (Azure Biosystems, Dublin, CA, USA).

#### Plasmid amplification, purification, and AAV packaging

Confirmed colonies were expanded in 2x Yeast Extract Tryptone (YT) media, and plasmid DNA was isolated using a Purelink HiPure Plasmid Filter Midiprep kit (Invitrogen, K210014) according to manufacturer’s instructions. Plasmid identity was confirmed by Oxford Nanopore plasmid sequencing (Plasmidsaurus, San Francisco, CA, USA). Purified plasmid DNA was subsequently used for in vitro validation of sgRNA activity and Cre-dependency (see below). The Cre-dependent constructs (pAAV-FLEX-SaCas9-U6-sg*Oprm1* and pAAV-FLEX-SaCas9-U6-sg*lacZ*) were then packaged into an AAV9 backbone by VectorBuilder (Chicago, IL, USA).

### In vitro validation of sgRNA

#### Nucleofection

We first validated sgRNA activity in vitro using nucleofected C6 rat glial cells (**Fig. S5 a, b**). Nucleofection was performed as previously described in Carullo et. al (2020) with modifications. C6 cells (American Type Culture Collection [CCL-107, ATCC, RRID:CVCL_0194]) were cultured in F-12k-based medium (Thermo Fisher Scientific, 21127022) supplemented with 12% Horse serum and 2.5% fetal bovine serum. Cells were seeded at a density of 95,000 cells/well in 12-well plates and allowed to reach 70-90% confluency over two days. Cells were washed with 1x sterile PBS, trypsinized (0.25% trypsin and 1 mM EDTA in PBS pH 7.4), and spun down at 900rpm for 5-7min. Cell pellets were then resuspended in nucleofection buffer (5 mM KCl, 15 mM MgCl, 15 mM HEPES, 125 mM Na_2_HPO_4_/NaH_2_PO_4_, 25 mM Mannitol) and nucleofected with 6.8 μg plasmid DNA per group. The Nucleofector™2b device (Lonza Bioscience, Walkersville, MD) was used according to manufacturer instructions (C6 high efficiency program U-030). For co-nucleofection of FLEX plasmids with mCherry-Cre or mCherry plasmids, DNA amounts were molar normalized to size of plasmids for a total of 6.8 μg. Nucleofection groups were diluted with 440 μl media respectively and plated in 12-well plates (∼312,500 cells/well) with four replicates per group. Plates underwent a full media change 6 hours after nucleofection and were imaged and frozen for downstream processing after 18 hours.

#### DNA Isolation, PCR, and sequencing

After nucleofection, genomic DNA was isolated from C6 cells using the DNeasy Mini Kit (Qiagen, 69504) according to manufacturer’s instructions with modifications for cultured cell samples. Briefly, samples were incubated with PBS master mix containing sterile 1x PBS, Proteinase K, and RNAse A (200 µL PBS, 20 µL Proteinase K, and 4 µL RNase A [100 mg/ mL] per sample) for 2 min at room temperature prior to lysis. Samples were then processed according to manufacturer’s protocol, including Buffer AL lysis, ethanol precipitation, column-based purification, and elution in Buffer EB. DNA was eluted in 30 µL Buffer EB.

PCR was used to amplify the genomic region surrounding the SaCas9 target cut site of *Oprm1*. Forward primer: 5’-CATGTTCACCAGCATATTCACC-3’; reverse primer: 5’-TCGGTTCAGACATCACCAATAC-3’. PCR was performed using Q5 High-Fidelity 2X Master Mix (New England Biolabs, M0492S), forward and reverse primers, genomic C6 DNA, and nuclease-free water on a C1000 Thermal Cycler (Bio Rad, Hercules, CA, USA) with the following thermocycler program: Initial denaturation for 30 seconds at 98ºC, 32 cycles of PCR (10 seconds at 98ºC for denaturation, 30 seconds at 65ºC for annealing, and 30 seconds at 72ºC for extension), and final extension of 2 minutes at 72ºC. Reaction products were then purified using a PCR Purification Kit (Qiagen, 28106). Following PCR purification, sampleswereoutsourcedto Plasmidsaurus (San Francisco, CA, USA) for Oxford Nanopore long-read sequencing, alignment to reference locus, and edit calling. Sequencing of the targeted locus following nucleofection with a constitutively expressing SaCas9 construct confirmed editing at the predicted *Oprm1* target site (**Fig. S5c, d**). Sequencing of the targeted locus following co-nucleofection with FLEX-SaCas9 constructs and either a Cre-mCherry or mCherry control plasmid confirmed Cre-dependent editing of *Oprm1* (**Fig. S5e, f**).

### Stereotaxic surgery

Naïve adult male and female heterozygous *Chst9*^*Cre-tdTomato*^ rats were anesthetized with 4 to 5% isoflurane and placed in a stereotactic apparatus (Kopf instruments, Tujunca, CA, USA). Rats were maintained at a surgical plane of anesthesia with 2 to 3% isoflurane, and respiratory rates were monitored throughout surgery and maintained between 35 and 55 respirations per minute. Surgical coordinates were determined using the Paxinos and Watson rat brain atlas (sixth edition), targeting the NAc shell bilaterally. Under aseptic conditions, guide holes were drilled using a size 14 carbide drill at anterior/posterior (AP) +1.7 mm, mediolateral (ML) ±1.3 mm, and the infusion needle was lowered at 9.7º to dorsal/ventral (DV) −7.6 mm relative to Bregma^108^. All infusions were made using a gastight 30-gauge stainless steel injection needle and 10μl syringe (Hamilton Company, Reno, NV, USA). Viral constructs (**Table S2**) were infused bilaterally at a rate of 0.25 μl/min using a syringe pump (Harvard Apparatus, Holliston, MA, USA), totaling 1 μl per hemisphere. Needles remained in place for at least 10 min following each infusion to allow for diffusion of the virus. Infusion needles were slowly retracted, guide holes were filled with sterile bone wax, and the surgical incision was closed with 6-0 nylon sutures (Delasco, NL166013F13M). At the end of surgery, rats were administered buprenorphine (0.03 mg/kg) and carprofen (5 mg/kg) for analgesia, and topical bacitracin (500 units) was applied to the incision site as a skin protectant and antimicrobial treatment. Animals were placed on a heating pad covered by a sterile drape to maintain body temperatures during surgery.

### Drugs

Fentanyl citrate (MWI Veterinary Supply, Catalog #045741) was dissolved in sterile 0.9% sodium chloride and injected intraperitoneally at the indicated doses for fentanyl CPP testing (15 µg/kg) and acute fentanyl exposure experiments (30 µg/kg). For electrophysiological recordings, D-APV (Tocris, 01-061-00), picrotoxin (Sigma-Aldrich, P1675), and SCH50911 (Tocris, 0984) were prepared as stock solutions in sterile water and diluted into artificial cerebrospinal fluid (aCSF) to final bath concentrations of 5 µM, 100 µM, and 1 µM, respectively. NBQX (Tocris, 0373) was prepared as a stock solution in dimethyl sulfoxide (DMSO; Invitrogen, D1234) and diluted into aCSF to a final concentration of 10 µM. DAMGO (Sigma-Aldrich, E7384) and naloxone (Tocris, 0599) were bath applied during recordings at a final concentration of 1 µM each.

### Locomotor testing

For comparison between genotypes, wildtype (n = 11), heterozygous (n = 22), and homozygous (n = 8) *Chst9*^*Cre-tdTomato*^ rats were placed in a 43 cm by 43 cm plexiglass locomotor activity chamber (Med Associates Inc., St. Albans, VT, USA) with opaque white wall covering and an open top, and locomotor activity was monitored for 20 minutes. For baseline locomotor activity, one day prior to CPP testing, female rats (n = 8 for *lacZ*, 9 for *Oprm1*) were placed in the same activity chambers, and baseline locomotor activity was monitored for 20 minutes.

### CPP testing

A biased CPP assay was performed as previously described^109,110^ with heterozygous *Chst9*^*Cre-tdTomato*^ rats using a three-chamber apparatus with guillotine-style doors (Med Associates Inc., Fairfax, VT, USA), composed of two conditioning chambers measuring ∼27 cm by 21 cm by 22 cm, separated by a central compartment measuring 12 cm by 21 cm by 22 cm. The first conditioning chamber had a wire mesh floor and white walls, and the second had stainless steel bar flooring. The central compartment had grey walls and a solid floor. All chambers had clear, perforated acrylic lids fixed with overhead lights and a 16-channel infrared controller (Med Associates Inc.) to track rodent movement and position. CPP testing began 4 weeks after infusion surgeries and was preceded by a baseline locomotion assay. During the pre-conditioning test, female rats (n = 8 for *lacZ*, 9 for *Oprm1*) were placed in the central compartment and the guillotine-style doors were opened to allow free movement between all three chambers for a duration of 30 min to determine individual chamber preferences. Preference was determined by the total time spent in each box. Days 2 through 5 were conditioning days; between 8 am and 11 am the rats were given an IP injection of saline and immediately placed in the initially preferred chamber for a 30-min test session. In the afternoons between 1 pm and 4 pm, rats were given an IP injection of fentanyl (15 µg/kg) and placed in the initially non-preferred chamber. Fentanyl sessions were conducted at least four hours after saline sessions. A post-conditioning test was performed on day 6 wherein animals were given free access to all three chambers as in the pretest and duration of time spent in each chamber was recorded.

### Whole-cell patch-clamp electrophysiology

To assess opioid sensitivity of Chst9-MSNs, whole-cell patch-clamp recordings were conducted as previously described^110,111^ in NAc slices from homozygous or heterozygous *Chst9*^*Cre-tdTomato*^ rats. 40-65 days old rats were briefly anesthetized with isoflurane and transcardially perfused with oxygenated ice cold recovery solution: 93 mM N-methyl-d-glucamine, 2.5 mM KCl, 1.2 mM NaH_2_PO_4_, 30 mM NaHCO_3_, 20 mM 4-(2-hydroxyethyl)-1-piperazineethanesulfonic acid (HEPES), 25 mM glucose, 4 mM sodium ascorbate, 2 mM thiourea, 3 mM sodium pyruvate, 10 mM MgSO_4_(H_2_O)_7_, 0.5 mM CaCl_2_(H_2_O)_2_, and HCl added until pH was 7.3 to 7.4 with an osmolarity of 300 to 310 mOsmol. Coronal slices (300 μm) were prepared on a vibratome (Leica, VT1200S) containing recovery solution and then transferred to a Brain Slice Keeper (Automate Scientific, Berkeley, CA, USA) containing a holding solution for at least 1 hour before recording: 92 mM NaCl, 2.5 mM KCl, 1.2 mM NaH_2_PO_4_, 30 mM NaHCO_3_, 20 mM HEPES, 25 mM glucose, 4 mM sodium ascorbate, 2 mM thiourea, 3 mM sodium pyruvate, 2 mM MgSO_4_(H_2_O)_7_, 2 mM CaCl_2_(H_2_O)_2_, and 2 M NaOH added until pH reached 7.3 to 7.4 and osmolarity was 300 to 310 mOsmol. Patch pipettes were pulled from 1.5-mm borosilicate glass capillaries (World Precision Instruments, Sarasota, FL, USA; 4-to 8-MOhm resistance). Pipettes were filled with K-gluconate intracellular solution for intrinsic experiments. K-gluconate composition was 120 mM K-gluconate, 6 mM KCl, 10 mM HEPES, 4 mM adenosine 5′-triphosphate–Mg, 0.3 mM guanosine 5′-triphosphate-Na, and 0.1 mM EGTA and titrated to a pH of ∼7.2 with KOH. During recordings, slices were transferred to a perfusion chamber and continuously perfused with artificial cerebrospinal fluid (aCSF) at 31ºC at a rate of 4 to 7 ml/min: 119 mM NaCl, 2.5 mM KCl, 1 mM NaH_2_PO_4_, 26 mM NaHCO_3_, 11 mM dextrose, 1.3 mM MgSO_4_(H_2_O)_7_, and mM 2.5 CaCl_2_(H_2_O)_2_. aCSF also contained the synaptic blockers D-APV (Tocris, 01-061-00; 5 μM), NBQX (Tocris, 0373; 10 μM), picrotoxin (Sigma-Aldrich, P1675; 100 μM), and SCH50911 (Tocris, 0984; 1 μM) to block NMDA, AMPA, GABA_A_ and GABA_B_ receptors, respectively.

All recordings were performed in the NAc shell in tdTomato-positive cells that were visualized using CellSens software (Olympus). All recordings were performed under continuous (gap free) conditions. Once patched, cells were voltage clamped at -75 mV. After switching to current clamp, cells were held at incremental current injections starting at 10 pA until consistent action potentials were observed. Once stable baseline firing was established (minimum 5 minutes), DAMGO (Sigma-Aldrich, E7384; 1 µM) was applied to the circulating aCSF to assess the effects of µOR activation on Chst9-MSN firing activity and membrane potential. Following a minimum of 5 minutes of µOR agonism, we applied naloxone (Tocris, 0599; 1 µM) into the circulating aCSF to determine µOR-mediated effects were rapidly reversed. Recordings in their entirety lasted ∼45 minutes. Changes in firing rate and membrane potential were monitored continuously. For all patch-clamp experiments, cells collected from the same animal were grouped together for analysis. Changes in action potential frequency and resting membrane potential were analyzed using pClamp11 (Clampfit, Axon Instruments). Series access and cell input resistance were monitored throughout recordings to ensure cells were only used if the access had not changed more than 10%. All drugs and reagents were obtained from Sigma-Aldrich unless specified otherwise.

### Immunohistochemistry

#### Tissue collection

Rats were transcardially perfused with 1x PBS followed by 4% PFA until body rigidity was confirmed. Brains were placed in 4% PFA overnight before being cryoprotected in 15% then 30% sucrose. Brains were sectioned at 30 μm using a Leica CM 1860 cryostat (Deer Park, IL, USA). Free floating sections were stored at -20ºC in cryoprotectant consisting of 50 mM sodium phosphate buffer (33.3 mM Na_2_HPO_4_, 16.7 mM NaH_2_PO_4_), 154 mM NaCl, 30% (w/v) sucrose, and 30% (v/v) ethylene glycol in distilled water.

#### Immunolabeling

*Chst9*^*Cre-tdTomato*^ rat validation, projection mapping, µOR knockout: Free-floating sections were thawed and washed twice in PBS, permeabilized for 30 min at room temperature in 0.25% Triton X-100, and blocked for 1 hour at room temperature in blocking buffer (10% normal goat serum, 0.5% bovine serum albumin, 0.05% Tween-20, 25mM glycine). Sections were incubated with primary antibodies (**Table S3**) for 48 hours at 4ºC, washed three times in PBS, and incubated with Alexa Fluor-conjugated secondary antibodies (**Table S3**) for 2 hours at room temperature. For HA immunolabeling, signal amplification was performed following primary antibody incubation. Sections were incubated with biotinylated secondary antibodies (Vector Biolabs, Malvern, PA; 1:200 dilution) in blocking solution for 2 hours, washed three times in PBS, and incubated with streptavidin-conjugated fluorophores for 2 hours. Sections were then washed three times in PBS, counterstained with DAPI (1:1000 in PBS) and mounted with ProLong Glass Antifade Mountant (P36984, Thermo Fisher Scientific, Waltham MA, USA).

Phosphorylated pyruvate dehydrogenase (pPDH): IHC was performed as previously described^47^. Free-floating sections were thawed and washed twice in PBS two times and then blocked for 1 hour at room temperature in PBS containing 5% goat serum and 0.3% Triton X-100. Sections were then incubated overnight at 4ºC with primary antibodies (**Table S3**) diluted in PBS containing 2% BSA and 0.3% Triton X-100. Sections were washed three times in P containing 0.3% Triton X-100 and incubated at room temperature for 1 hour with secondary antibodies (**Table S3**) diluted in PBS containing 2% BSA and 0.3% Triton X-100. Sections were then washed three times, counterstained with DAPI (1:1000 in PBS), and mounted with ProLong Glass Antifade Mountant (P36984, Thermo Fisher Scientific, Waltham MA, USA).

#### Image quantification

Section images were acquired using the BZ-X800 Keyence Microscope and stitched using Keyence Image Analyzer software. 20x magnification images were taken at identical exposures per channel. Mean fluorescence intensity (MFI) was quantified in ImageJ software^112^. Regions of interest (ROI) were defined by HA immunoreactivity for µOR knockout validation or RFP immunoreactivity for pPDH analyses and applied to the corresponding µOR or pPDH channel. Background-corrected MFI was calculated by subtracting fluorescence measured in the anterior commissure from each ROI. For each animal, area-weighted MFI was calculated by dividing the summed area-weighted fluorescence by the total ROI area.

### Statistical Analysis

All statistical and graphical analyses were performed with Prism software (GraphPad, La Jolla, CA). Electrophysiological measurements of action potential frequency and resting membrane potential were analyzed using one-way repeated measures ANOVA with Tukey’s multiple comparisons tests. For fentanyl CPP, data were analyzed using a two-way repeated measures ANOVA with Šídák multiple comparisons correction. sgRNA treatment was included as between-subjects factor and time (pre-test vs post-test) as a within-subjects factor. Bodyweight and locomotor activity across genotypes were analyzed with ordinary one-way ANOVAs. Differences in pPDH immunoreactivity, µOR immunoreactivity, open-field behavior, and CPP difference scores were assessed using unpaired t-tests. Data are presented as mean ± SEM, and statistical significance was defined as *p* < 0.05.

Potential sex-dependent effects of fentanyl on pPDH immunoreactivity were assessed using a three-way mixed-effects ANOVA on raw (non-normalized) pPDH immunoreactivity measurements with treatment and sex as between-subjects factors and cell type (tdTomato+ vs tdTomato-) as within-subjects factor. This analysis revealed a significant Treatment x Cell type interaction (F_(1,17)_ = 9.87, *p* = 0.004), indicating differential effects of fentanyl across cell populations. The Treatment x Sex x Cell type interaction was not significant (F_(1,17)_ = 2.03, *p* = 0.167), indicating that the cell-type-specific response to fentanyl was not significantly influenced by sex. Because sex did not significantly influence fentanyl-induced changes in pPDH immunoreactivity, data from male and female rats were combined.

## Data availability

Code for analyses of snRNA-seq data is available at https://github.com/Jeremy-Day-Lab/Drake_2026_NAc_snRNAseq, and both raw and processed data files supporting this experiment are available in Gene Expression Omnibus (GSE337210). Single nucleus RNA sequencing data can also be freely viewed at www.ratlas.org. Code for analysis of the Xenium spatial transcriptomics data is available at https://github.com/Jeremy-Day-Lab/Drake_2026_Xenium_AP, and both raw and processed data files supporting this experiment are available in Gene Expression Omnibus (GSE337270). R objects for snRNA-seq and Xenium datasets can be found at doi: 10.5281/ zenodo.21109895.

## ACKNOWLEDGEMENTS

We thank all Day Lab members for assistance and support. We acknowledge the UAB Flow Cytometry & Single Cell Core Facility, the UAB Genomics Core Laboratory, the UAB Biological Data Sciences Core (RRID:SCR_0217666), the UAB High Resolution Imaging Facility, and the UAB Animal Behavioral Assessment Core.

## FUNDING

NIH grants R01DA053743, R01DA054714, and R01DA066682 (JJD)

UAB MPI Award (JJD, JAH, JP)

T32 MH129274 (ORD)

## AUTHOR CONTRIBUTIONS

Conceptualization: ORD, JP, JAH, JJD

Methodology: ORD, CMF, ETJ, CEN, AP, JJD

Investigation: ORD, CMF, ETJ, CEN, AP, JJD

Formal analysis: ORD, CMF, ETJ, CEN, LAP, AT, LI, & JJD

Software: ORD, CEN, LAP, AT, LI, JJD

Resources: ORD, ETJ, LI, JP, JAH, & JJD

Funding acquisition: ORD, JP, JAH, & JJD

Data curation: ORD, CMF, CEN, LAP, AT, AP, JJD

Visualization: ORD, CMF, CEN, JJD

Validation: ORD, CMF, JJD

Project administration: ORD, JP, JAH, JJD

Supervision: LI, JP, JAH, JJD

Writing—original draft: ORD, CMF, CEN, JJD

Writing—review & editing: ORD, CMF, ETJ, CEN, LAP, AT, AP, LI, JP, JAH, & JJD

---

**Supplementary Figure 1.**
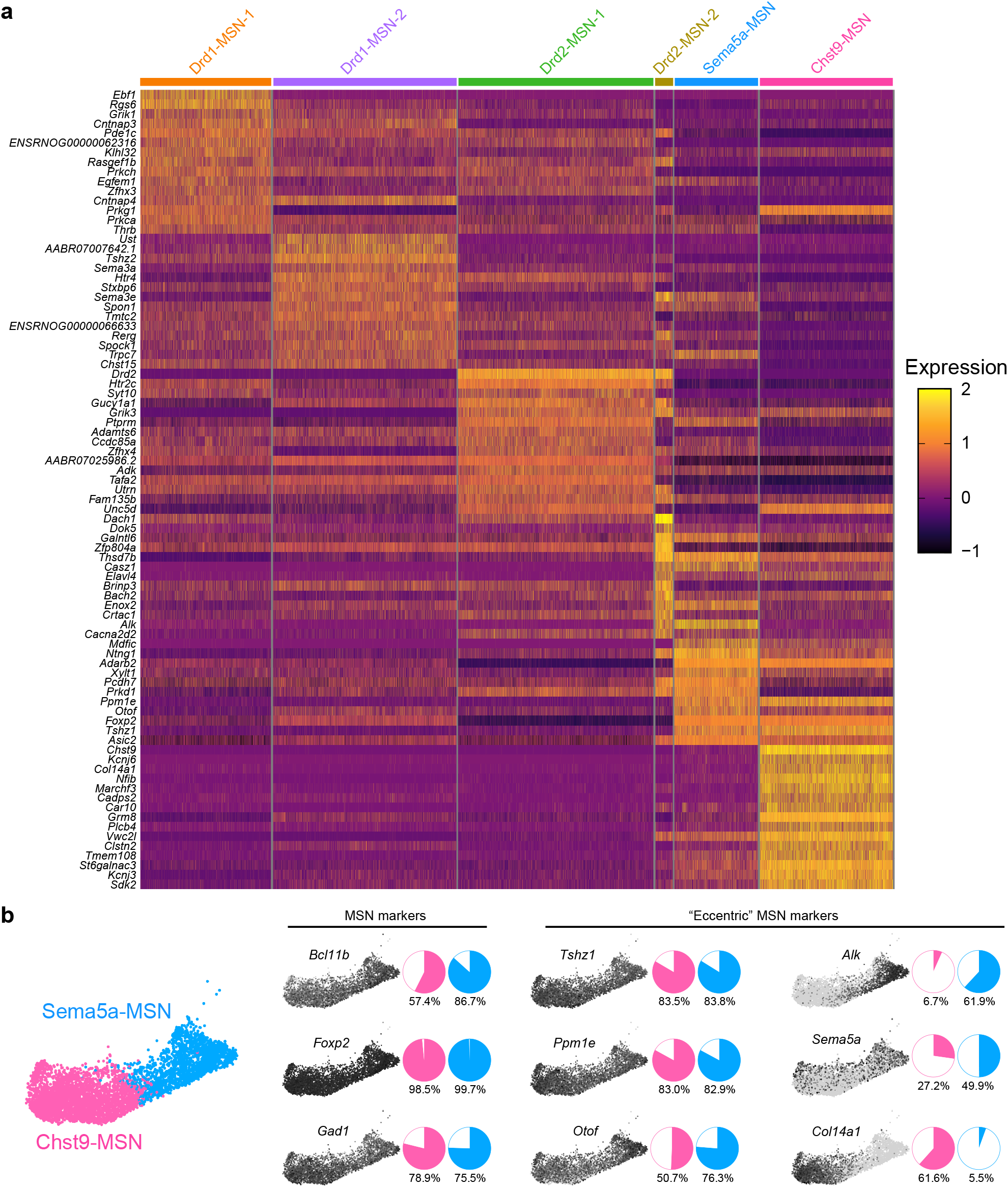
Differential gene expression among medium spiny neuron (MSN) populations. **a**, Heatmap showing differentially expressed genes (DEGs) across MSN populations. Genes were filtered to include the top 15 DEGs per population (adjusted p value < 0.05, average log_2_ fold change > 0.5, and >50% of cells expressing). **b**, UMAP of Chst9-MSN and Sema5a-MSN populations (left). Feature plots and quantification of the percentage of cells expressing canonical MSN markers (middle) and previously identified markers associated with “eccentric” MSN populations (right).

**Supplementary Figure 2.**
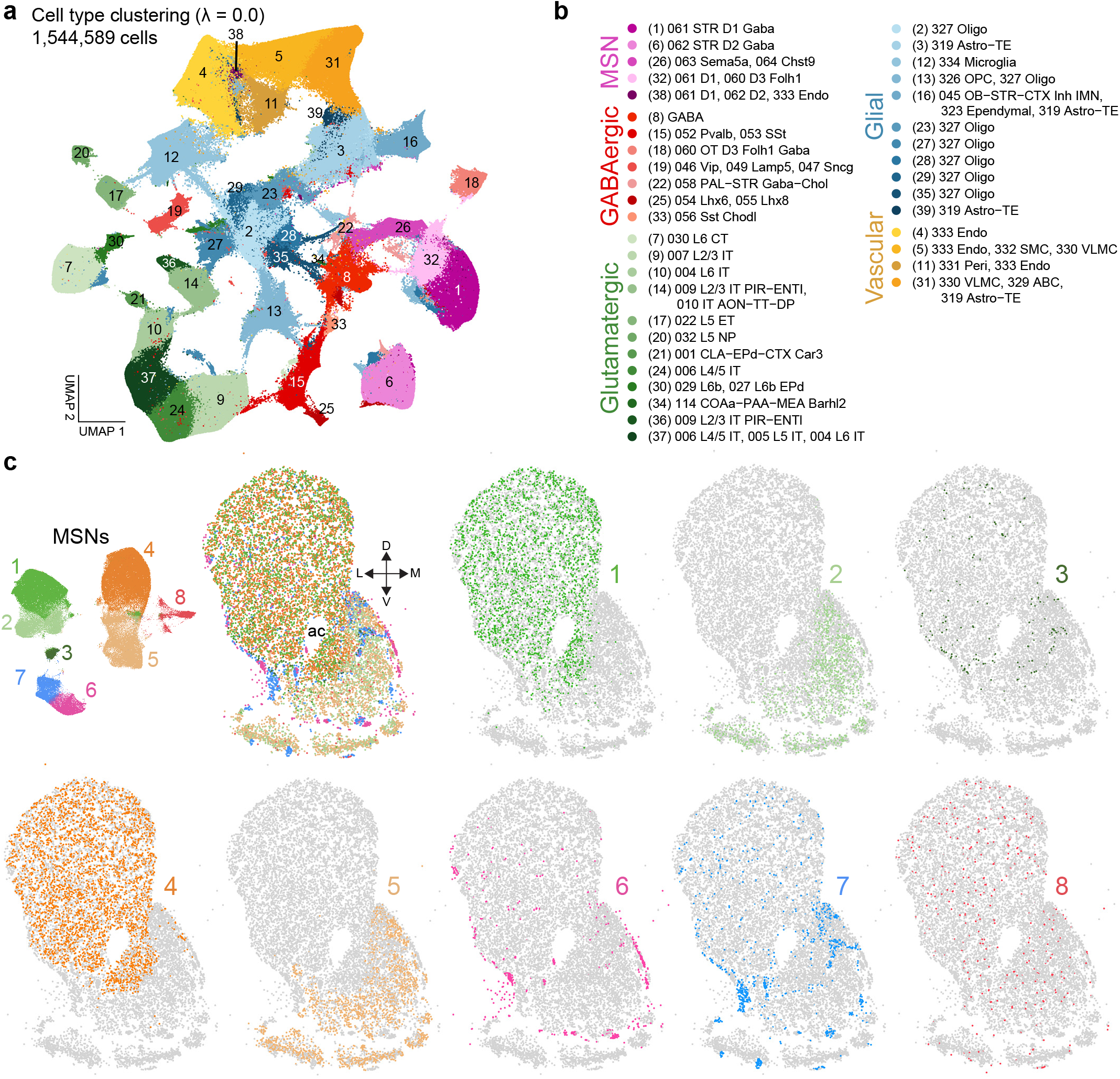
Cell type clustering of Xenium spatial transcriptomics data identifies MSN subclasses in the nucleus accumbens. **a**, Unsupervised clustering of Xenium spatial transcriptomics data was performed using 30 principal components and a clustering resolution of 1.0. UMAP visualization of 1,544,589 cells across 16 coronal sections collected from 2 male and 2 female mice. **b**, Cell populations annotated using the Allen Institute Whole Mouse Brain taxonomy and colored by major cell class, including medium spiny neurons (MSNs), GABAergic interneurons, glutamatergic neurons, glial cells, and vascular-associated cells. These annotations were used to identify MSN populations for downstream analyses. **c**, UMAP visualization of MSN populations following subsetting and reclustering using the same clustering parameters (top left). Representative spatial maps from a single animal showing the distribution of all MSNs together and individual spatial distributions of each MSN subtype.

**Supplementary Figure 3.**
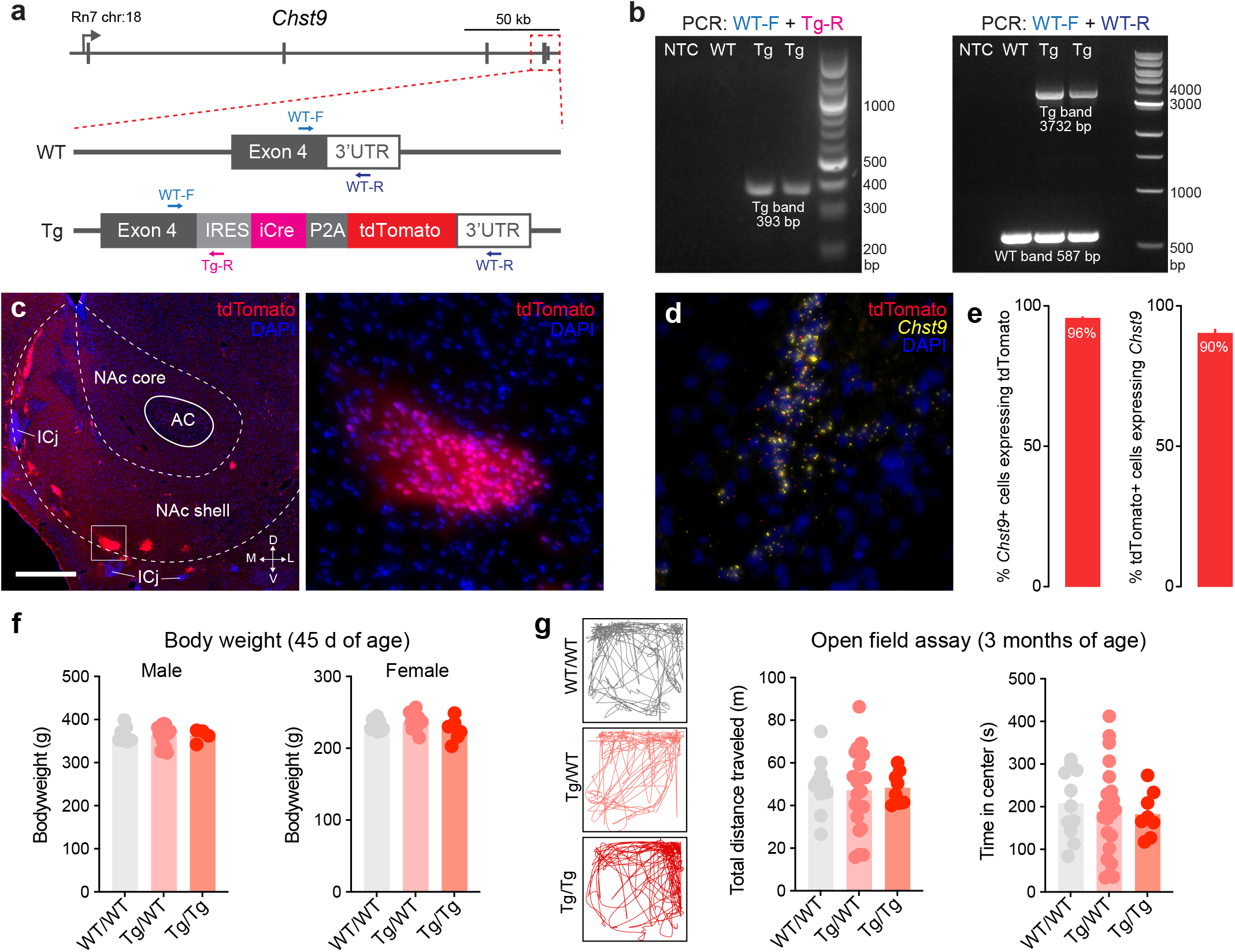
Generation & validation of a Sprague-Dawley-*Chst9*^*IRES-iCre-P2A-tdTomato*^ (*Chst9*^*Cre-tdTomato*^) transgenic rat. **a**, Location of IRES-NLS-iCre-P2A-tdTomato cassette inserted following exon 4 of the rat *Chst9* gene. Insertion was performed using CRISPR-Cas9 mediated homology directed repair. PCR primers used for validation are shown in bottom panel. **b**, Accurate transgene insertion in heterozygous founders was confirmed by PCR of genomic DNA. Left, heterozygous transgenics harbor a single 393 bp PCR product when using WT-F and Tg-R primer pair. Right, PCR amplification using WT-F and WT-R primers results in a 587 bp product in both wild type and heterozygous *Chst9*^*Cre-tdTomato*^ rats, but only transgenic rats harbor a larger 3732 bp band corresponding to the transgene insert size. NTC = no template control. **c**, Left, tdTomato fluorescence (detected with IHC) in the NAc of a Chst9-IRES-iCre-P2A-tdTomato transgenic rat. ICj = Islands of Calleja. Right, 100x magnification image of Chst9-MSN cluster. **d**, Single molecule RNA fluorescence in situ hybridization (smRNA FISH) validation of overlap between *Chst9* and tdTomato mRNA. **e**, smRNA FISH quantification verifies selective expression of tdTomato in *Chst9*+ cells (n=3). Similar values are obtained for overlap between *Chst9* and Cre mRNA. **f**, Genotype had no effect on body weight in males (left, Ordinary one-way ANOVA, F_(2,27)_ = 0.001182, *p* = 0.9988) or females (right; Ordinary one-way ANOVA, F_(2,26)_ = 1.795, *p* = 0.1862). **g**, Open field test. Left, representative traces showing animal location during 20 minute test session. Genotype did not affect total distance traveled (middle; Ordinary one-way ANOVA, F_(2,38)_ = 0.08069, *p* = 0.9226.) or time spent in the center of field (right; Ordinary one-way ANOVA, F_(2,38)_ = 0.2148, *p* = 0.8077).

**Supplementary Figure 4.**
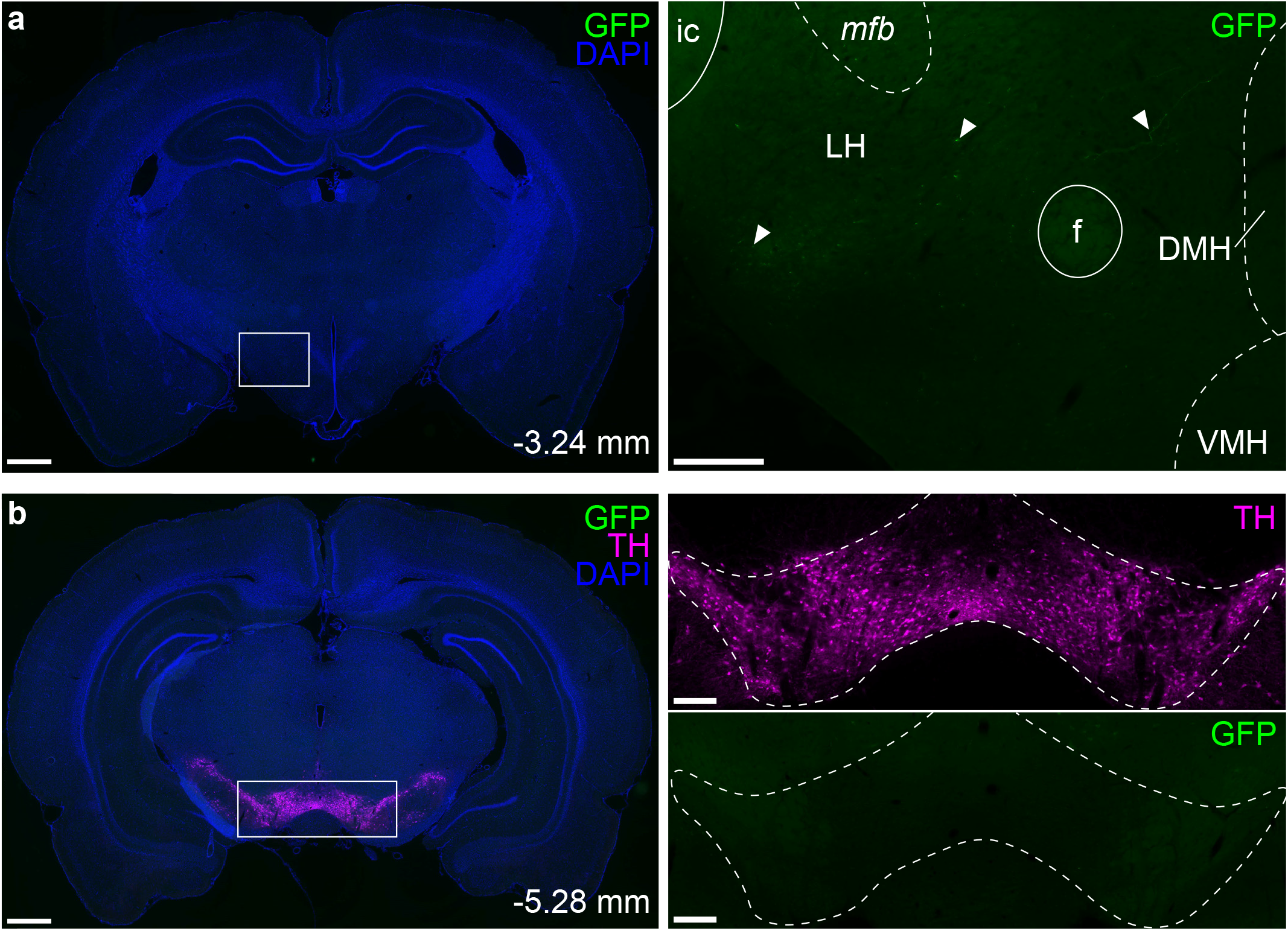
GFP labeling in the lateral hypothalamus and ventral tegmental area following Chst9-MSN tracing. **a**, Representative coronal section containing the lateral hypothalamus (LH) following injection of a Cre-dependent GFP reporter into the nucleus accumbens of *Chst9*^*Cre-tdTomato*^ rats. Left, 4x image showing GFP (green) and DAPI (blue); scale bar = 1000 µm. Right, 20x image of the LH showing sparse GFP-positive fibers. ic, internal capsule; mfb, medial forebrain bundle; f, fornix; DMH, dorsomedial hypothalamus; VMH, ventromedial hypothalamus; scale bar = 250 µm. **b**, Representative coronal section containing the ventral tegmental area (VTA). Left, 4x image showing GFP (green), tyrosine hydroxylase (TH, magenta), and DAPI (blue); scale bar = 1000 µm. Right, 20x images showing TH immunostaining delineating the VTA (magenta) and corresponding GFP expression (green). Dashed lines indicate the approximate boundaries of the VTA; scale bar = 250 µm.

**Supplementary Figure 5.**
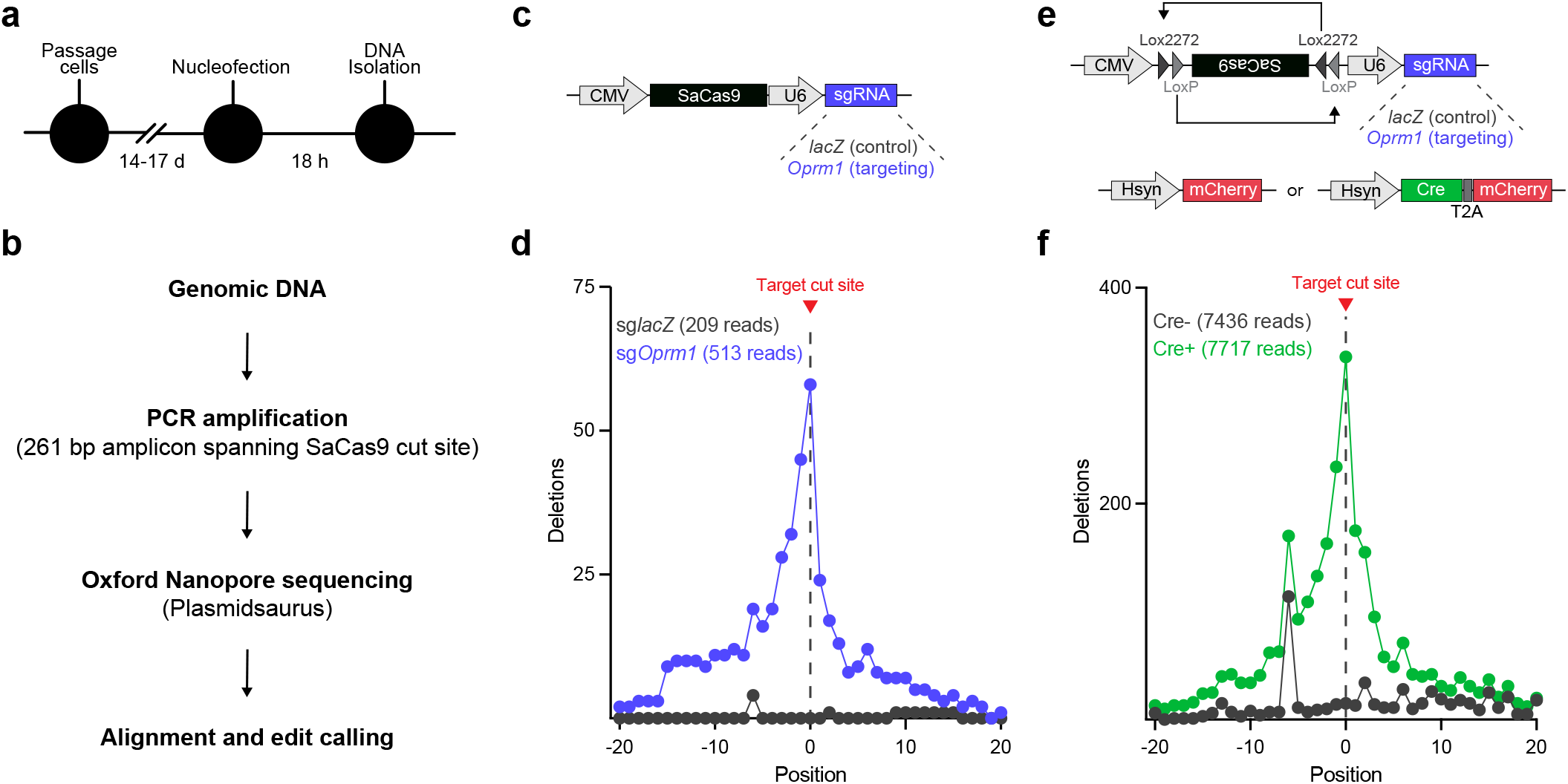
In vitro validation of *Oprm1 s*gRNA. **a**, Nucleofection strategy. Cells were passaged and nucleofected on passage days 14-17, followed by DNA isolation 18 h later. **b**, Genomic DNA was PCR amplified across the target cut site, subjected to long-read sequencing, aligned to reference locus, and analyzed for editing outcomes. **c**, Cells were nucleofected with a plasmid constitutively expressing SaCas9 and sgRNAs. **d**, Number of deletions surrounding the sgRNA target site in cells nucleofected with the constitutively expressing *Oprm1* plasmid compared to *lacZ* control. **e**, Cells were co-nucleofected with FLEX-SaCas9 constructs and either a Cre-mCherry plasmid or an mCherry control plasmid. **f**, Number of deletions surrounding the sgRNA target site in cells nucleofected with FLEX-SaCas9 plasmids with or without co-nucleofection of Cre.

